# Gap-free genomes and transcriptomes uncover race-specific effectors in watermelon wilt pathogen *Fusarium oxysporum* f. sp. *niveum*

**DOI:** 10.1101/2025.04.04.647254

**Authors:** Dilay Hazal Ayhan, Huan Wang, Lili Zhang, Guan Wang, Shu Yi, Dian Meng, Lifang Xue, Xin Geng, Zhijun Kong, Xinrui Wang, Lu Wang, Qingxian Yang, Xiangfeng Wang, Yun Deng, Xingping Zhang, Li Guo

## Abstract

Watermelon (*Citrullus lanatus* L.) is one of the world’s most economically important fruit crops, yet it is susceptible to devastating diseases such as vascular wilt caused by *Fusarium oxysporum* f. sp. *niveum* (Fon), with limited effective control measures. Fon evolves rapidly in the field to overcome host resistance and global watermelon production is constantly threatened by new pathogenic races. High-quality genomic resources are key to understanding the molecular mechanisms underlying Fon virulence evolution for disease management. Here, we *de novo* assembled and annotated gapless genomes of three physiological races of Fon (race 1, 2, and 3), and dissected the mechanisms behind their distinctive virulence through comparative genomics and transcriptomics. Whole-genome alignments identified core and accessory chromosomes in Fon where each race carried a unique set of accessory chromosomes or regions. Comparative transcriptomics of Fon infection revealed distinctive temporal patterns of gene expression even among core gene families, particularly those related to cell wall degradation enzymes. Effectoromic prediction and comparative analysis in three gap-free genomes identified 13 race-specific effectors (RSEs) in FonR3, one (FonRSE1) of which was a critical virulence factor of FonR3 on watermelon as demonstrated via functional experiments. The gap-free genome assemblies and annotations, and the RSEs are valuable resources for studying Fon pathobiology and genome evolution, adding in the design of improved disease control strategies.

## INTRODUCTION

*Fusarium oxysporum* is a widely spread phytopathogenic fungus causing Fusarium wilt in over 120 plant species^1^ and considered among the ten most important fungal plant pathogens^2^. The soilborne fungus invades the plant vascular system with mycelium through roots and causes embolism, leading to plant wilting and eventual death. Despite the broad host range of *F. oxysporum* species complex (FOSC), an individual strain of *F. oxysporum* is host-specific^3^, and strains virulent to the same host range are grouped into formae speciales (f. sp.). Fusarium wilt of watermelon (*Citrullus lanatus* L.) caused by *F. oxysporum* f. sp. *niveum* (Fon) is often regarded as the most devastating disease of watermelon, one of the most important fruit crops with great nutritional and economic values worldwide^4^. Although several watermelon germplasms confer resistance to Fusarium vascular wilt^5^, the evolution of Fon quickly overcomes the resistance giving rise to various physiological races. So far, four races (race 0, 1, 2, and 3) of Fon have been recognized based on pathogenicity test using watermelon differential cultivars^6–9^. Among the four races, Fon race 0 is the least economically important being non-pathogenic to the majority of commercial cultivars^4^. Watermelon cultivars such as Calhoun Gray and Sugarlee are resistant to Fon race 1. However, such resistance was overcome by Fon race 2 and no resistant resource was available to Fon race 2 until the release of cultivar PI-296341-*FR* and PKR6^5,10,11^. Unfortunately, Fon race 3, according to recent reports (USA and China), can overcome all tested watermelon cultivars, including the highly resistant PI-296341-*FR* and PKR6^8,9,12^. Thus, Fon race 3 poses a huge risk for global watermelon production, requiring research and quarantine efforts to contain its dissemination.

Genomics is essential to understanding the pathogen evolution and dissecting the molecular mechanisms underlying the fungal virulence. *F. oxysporum* genomes, pathogenic or not, are typically organized into two compartments, the core chromosomes (CCs) and accessory chromosomes (ACs)^3^. CCs are genomically conserved among *F. oxysporum*, gene-rich, and vertically transmitted, whereas ACs are typically repeat-rich, lineage- or strain-specific, and could be horizontally transferred among strains. *F. oxysporum* generally has 11 CCs which encode genes essential to survival and reproduction, whereas the number of ACs is highly variable among different formae specialis or even races, and AC genes are associated with the fungal pathogenicity and host specificity of the strain. Furthermore, ACs have distinctive characteristics in genomic compositions than CCs such as gene-sparse, low GC content, high transposable element (TE) density, and H3K27me3 methylation indicative of heterochromatin status^13^, presenting criteria to identify the ACs in *F. oxysporum* genomes. Genomic analysis of *F. oxysporum* has shown that ACs are enriched in effectors as virulence factors helping pathogens in colonization and infection of host cells. Effectors are small secreted proteins carrying signal peptides, rich in cysteine residues and induced by host infection^14,15^. After the secretion into plant-pathogen interfaces, effectors can either kill host cells with toxicity or bind with the target proteins of host plants to inhibit their immune responses and manipulate host cell activities^16,17^. Interestingly, effector genes are usually arranged in flexible genomic regions with high TE contents, which might aid the rapid evolution and loss-and-gain of virulence and host specificity^16^. Therefore, effectors are potentially key players in Fon virulence to watermelon and the different repertoire of effectors in Fon races could account for the race evolution. Effector studies in Fon have been limited so far, requiring high-quality reference genomes. There have been published genome assemblies for different Fon races^18^. However, the published Fon genomes are draft assemblies at the contig-level, containing many gaps, given the highly repetitive nature of *F. oxysporum* genomes, which are notoriously difficult to assemble. Currently, gap-free Fon genomes have not been reported, and there have not yet been in-depth comparative genomics and transcriptomics studies among various races to understand the genetic basis and virulence evolution.

In this study, we assembled and annotated gap-free genomes for three isolates representing Fon races 1, 2, and 3, and performed comparative genomics and transcriptomics to identify putative determinants of virulence differentiation in Fon races. Genome comparison identified CCs and ACs from three races and revealed conserved and unique accessory regions and chromosomes. Comparative transcriptomics uncovered different temporal patterns of gene expression in three races, even for orthologous genes especially related to cell-wall degradation. We further identified FonR3 (race 3) specific effectors essential for its full virulence on previously resistant PKR6. Our study provided valuable and high-quality genomic resources as well as effector profiles for studying the virulence mechanism of Fon. The results also shed light on the importance of race-specific effectors on the emergence of pathogenic races and facilitate breeding resistant watermelon cultivars.

## RESULTS

### Pathogenicity and phylogenetics of three Fon physiological races

For this study, we obtained three Fon strains (Fon-1, Fon-2 and Fon-3) isolated on diseased watermelons from Crop Genetics and Breeding Platform of Peking University Institute of Advanced Agricultural Sciences. To verify their pathogenicity to cause Fusarium wilt of watermelons, the three Fon strains were inoculated against a set of watermelon differential cultivars used in pathogen race identification including G42 (susceptible to Fon race 1, 2, and 3), Calhoun Gray and Sugarlee (susceptible to Fon race 2 and 3 but resistant to race 1), and PKR6 (susceptible to Fon race 3 but resistant to race 1 and 2)^19,5,8,11^. Pyramiding multiple disease resistance, PKR6 is a newly released inbred line conferring high-level resistance to Fon race 2 but not 3, and thus chosen as a key differential host to identify race 3 and investigate the specific effector repertoire of race 3^11^. Our greenhouse bioassay results showed that Fon-1 was pathogenic only to G42, leading to 100% seedling death of G42 (**Fig. 1A**). Fon-2 overcame G42 and Calhoun Gray completely, and caused 80% death incidence on Sugarlee, but caused no detectable symptoms at PKR6 seedlings (**Fig. 1A**). In contrast, Fon-3 caused vascular wilt symptoms such as stunting, yellowing, wilting or plant death on seedlings of all four tested cultivars (**Fig. 1A**). Therefore, based on the disease symptoms of watermelon differential hosts, Fon-1, Fon-2, and Fon-3 were verified to be Fon race 1, 2, and 3, named FonR1, FonR2 and FonR3, respectively. To understand the evolutionary relationship of the three races with other FOSC members, we constructed a phylogenetic tree including three Fon isolates and 17 other *F. oxysporum* strains rooted with *F. verticillioides* strain BRIP53590 single-copy orthologous genes (see Methods) and annotated the clades as described in O’Donnell et al.^20^ (**Fig. 1B**). All three Fon isolates were located in Clade 2, where FonR1 and FonR3 were grouped in one subclade and phylogenetically close to each other, but both were distant from FonR2, located in another subclade. Two subclades containing Fon isolates also harbored other formae speciales (**Fig. 1B**). This was consistent with a previous phylogeny analysis revealing that Fon and other members of FOSC were generally polyphyletic and dispersed in distinct clades or subclades^16^. This reflected a complex evolutionary route for Fon races. To understand the genomic basis underlying virulence differentiation, the three isolates representing three pathogenic races of Fon were subjected to downstream whole-genome sequencing, assembly, annotation, and comparative analysis.

**Fig. 1.**
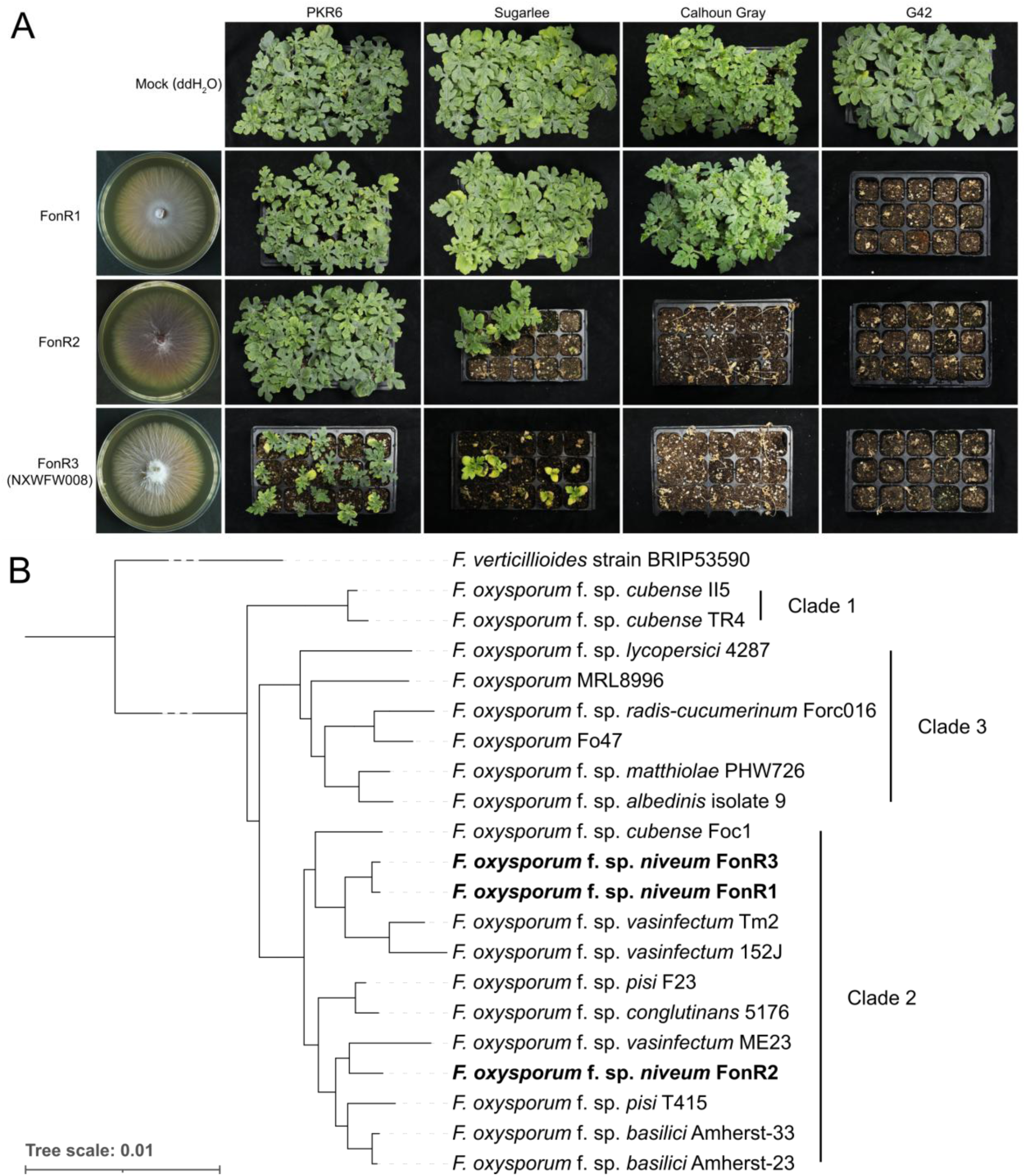
Phenotypes and phylogeny of three races of *Fusarium oxysporum* f. sp. *niveum* (Fon) sequenced in this study. (A) Potato dextrose agar (PDA) cultures of FonR1, FonR2, and FonR3 (NXWFW008) and disease phenotypes on differential hosts. PDA cultures were photographed 5 days post inoculation (dpi), while the plant infections were 28 dpi on 11-day-old seedlings. Mock was inoculated with water. Different watermelon plants used to identify the physiological races of Fon were PKR6 (resistant to R1, R2, but susceptible to R3), Sugarlee and Calhoun Gray (resistant to R1, but susceptible to R2 and R3), and G42 (susceptible to all three races). For each treatment, 15 plants were inoculated, and the experiment was repeated three times. (B) Phylogenetic species tree of 20 *F. oxysporum* strains including FonR1, FonR2, and FonR3. The tree was rooted by *F. verticillioides* strain BRIP53590. All branches are supported by 100% bootstrap values. Clades were annotated as described in O’Donnell et al.^20^.

### Gapless genome assemblies and annotations of three Fon races

To understand the evolution of pathogenicity in Fon races at the genomic level, we *de novo* assembled high-quality gapless reference genomes of FonR1, FonR2, and FonR3, which will reveal how the three races differ at the chromosome level. We first generated high-coverage sequencing data from each race using a combination of sequencing technologies, including PacBio high-fidelity (HiFi) long reads, Oxford Nanopore Technology (ONT) ultra-long reads, and high-throughput chromatin conformation capture (Hi-C) sequencing reads. Three draft genomes were assembled from HiFi and ONT ultra-long reads using Hifiasm^21^, followed by scaffolding to pseudochromosomes using Hi-C reads using the Juicer/3D-DNA pipeline^22,23^. The Hi-C contact maps indicated 18, 13, and 16 pseudochromosomes in the FonR1, FonR2, and FonR3 genome assemblies, respectively (**Fig. S1**). The ONT reads were applied to fill the remaining gaps in the initial assemblies using TGS-gapcloser^24^, obtaining three gapless genome assemblies (**Fig. S2**). The chromosome IDs of the assemblies were sorted in order of decreasing chromosome length. The assemblies had 64.66, 57.66, and 61.36 Mb genome sizes and contig N50 lengths of 3.72, 4.34, and 4.34 Mb for FonR1, FonR2, and FonR3, respectively (**Table 1**). All centromeres and 41.5% of the telomeres (20/36, 17/26, and 2/32 in FonR1, FonR2, and FonR3, respectively) were successfully captured (**Fig. 2A; Fig. S3**). To validate the assemblies, we mapped raw sequencing reads to each assembly, showing a high mapping rate of HiFi (98.72%-99.79%), ONT (99.82%-100%) and Illumina (92.20%-94.60%) reads. In addition, the Benchmarking Universal Single-Copy Ortholog (BUSCO) assessment^25^ resulted in 99.4%, 99.5%, and 99.4% completeness with QV values of 46.4, 57.29, and 40.36 for FonR1, FonR2, and FonR3, respectively (**Table 1**). Together, these indicate high accuracy and completeness of the three gapless genome assemblies.

**Fig. 2.**
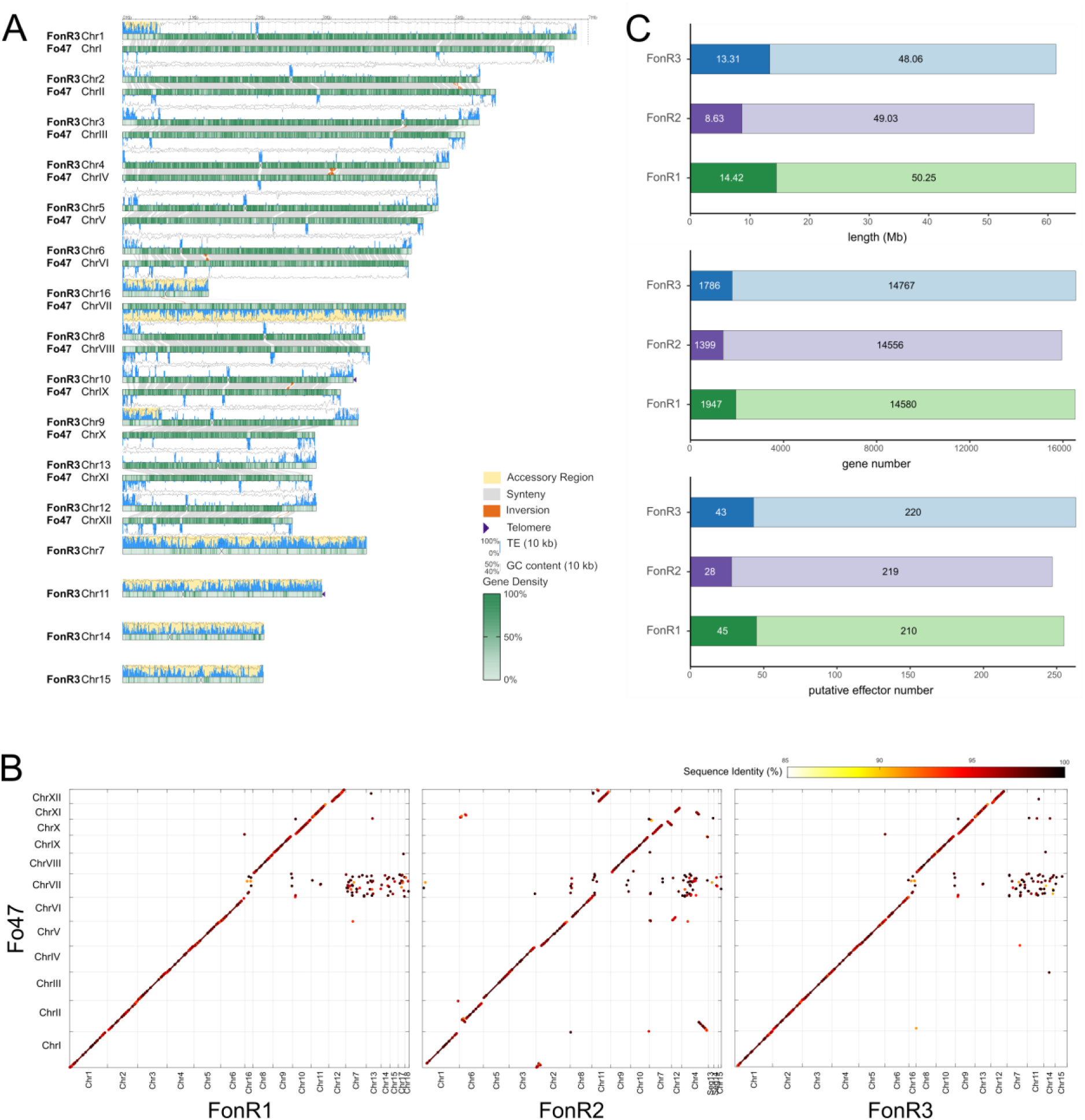
Gap-free genome assembly of Fon race 3 and accessory genome identification of FonR1, FonR2, FonR3. (A) An ideogram of the most virulent race, FonR3, compared to the Fo47 genome was visualized using GenomeSyn^77^. Genomic features, including gene density, GC content, TE content, telomeres, centromeres, and genome synteny with Fo47 chromosomes, are shown on the ideograms. The yellow background indicates the accessory regions. (B) Dotplots of FonR1, FonR2, and FonR3 genome alignments with Fo47. The color scale shows the sequence identity of the aligned sequences. (C) Comparison of three races in terms of total length, the number of annotated genes, and predicted effectors in core (light colors) and accessory (dark colors) genomic regions.

**Table 1.** Genome assembly and annotation statistics for three races of *Fusarium oxysporum* f. sp. *niveum*.

| Statistics | FonR1 | FonR2 | FonR3 |
| --- | --- | --- | --- |
| Genome size (bp) | 64,668,425 | 57,664,902 | 61,367,986 |
| Chromosome number | 18 | 13 | 16 |
| Contigs > 500 kb | 18 | 15 | 16 |
| Contigs > 50 kb | 45 | 49 | 19 |
| Contig number | 75 | 82 | 86 |
| Contig N50 (bp) | 3,727,044 | 4,345,875 | 4,344,065 |
| Contig N90 (bp) | 1,413,554 | 1,004,243 | 2,123,745 |
| Gap number | 0 | 0 | 0 |
| GC content (%) | 47.83 | 48.09 | 47.67 |
| TE content (%) | 20.35 | 18.51 | 16.82 |
| BUSCO completeness (%) | 99.4 | 99.5 | 99.4 |
| QV | 46.14 | 57.29 | 40.36 |
| Gene number | 16,527 | 15,955 | 16,553 |
| BUSCO completeness for genes (%) | 97.1 | 99.5 | 97.5 |
| TE content (%) | 20.35 | 18.51 | 16.82 |
| TE length (bp) | 13,160,780 | 10,675,578 | 10,324,453 |
| Retroelements (%) | 4.13 | 3.51 | 4.23 |
| LTR elements (%) | 2.58 | 2.23 | 2.56 |
| DNA transposons (%) | 6.01 | 5.7 | 5.84 |
| Putative effector number | 255 | 247 | 263 |

We then annotated the three gapless genomes by integrating *ab initio* prediction, homology, and transcriptome evidence using the MAKER pipeline^26^. To ensure accurate annotation of genes involved in fungal virulence, we sequenced transcriptomes (RNA-seq) of watermelon plants infected by each of three races at 1, 3, and 6 days post inoculation (dpi) using Illumina paired-end sequencing, with fungal mycelia harvested from PDB culture (0 dpi) as control. The Fon genome assemblies were first masked for the repeat sequences identified through RepeatMasker^27^, followed by protein-coding gene calling. A total of 16,527, 15,955, and 16,553 protein-coding gene models were predicted for FonR1, FonR2, and FonR3, respectively (**Table 1**), 36.94%, 37.67%, and 37.23% of which can be functionally annotated by either Gene Ontology (GO) or Kyoto Encyclopedia of Genes and Genomes (KEGG). The comparison of the protein-coding genes using OrthoFinder suggested that 391, 1113, and 464 were race-specific genes in FonR1, FonR2, and FonR3, respectively. The repeat contents detected by RepeatMasker in whole genomes for FonR1, FonR2 and FonR3 were 20.35%, 18.51%, and 16.82% where the retroelement contents were 4.13%, 3.51%, and 4.23%, and the DNA transposon contents were 6.01%, 5.70%, and 5.84%, respectively (**Table 1**).

### Structure of Fon core and accessory chromosomes/regions

*F. oxysporum* genomes are well known to have two major compartments: core and accessory genomes, with the latter able to horizontally transfer among different strains and thus contribute to the evolution of pathogenicity^3^. To identify core and accessory genomes for the three new assemblies, we performed whole-genome sequence alignment of FonR1, FonR2, and FonR3 assemblies against the complete genome of a nonpathogenic *F. oxysporum* strain Fo47 (NCBI Accession: GCA_013085055.1)^28^ which has 11 core and one accessory chromosomes (**Fig. 2A, B; Fig. S3**). Overall, the 11 core chromosomes of Fo47 were overall well aligned with 11 chromosomes in each Fon race despite a low alignment rate for repeat-rich regions, indicating that these were Fon core chromosomes (**Fig. 2A; Fig. S3**). We then defined the Fon accessory genomes as genomic regions longer than 400 kb, unaligned with Fo47 core chromosomes, and displaying the genomic characteristics of *F. oxysporum* accessory chromosomes, *e.g.,* low gene density and high repeat content. Under such stringent criteria, 14.42, 8.63, and 13.31 Mb sequences representing accessory genomes were identified in FonR1, FonR2, and FonR3, respectively (**Fig. 2C**). Specifically, Chr07, Chr13-18 in FonR1, Chr15 in FonR2, Chr07, Chr11, Chr14-16 in FonR3 were assigned as accessory chromosomes (**Fig. 2A; Fig. S3**, yellow background). In addition, parts of Chr01, Chr10, and Chr11 in FonR1, unmapped segments Chr13 and Chr14 and parts of Chr 01, Chr02, Chr04, Chr07, and Chr11 in FonR2, and parts of Chr01, Chr09 in FonR3 were identified as accessory regions. Interestingly, accessory chromosomes or regions in FonR1 and FonR3 were mostly standalone without attaching to other chromosomes, while accessory genomes in FonR2 mainly exist as "accessory/core chimera chromosomes" likely resulted from large-scale genome rearrangement in FonR2 (also visible in **Fig. 2B**). Notably, unlike other accessory regions attached to ends of core chromosomes, a particular accessory region (2.27-3.38 Mb) of FonR2 resided in the middle of the chromosome (Chr07) surrounded by two core regions (**Fig. S3**), an observation that has not been reported in *F. oxysporum* genomes so far. The core-accessory boundaries were confirmed by both long-read alignments spanning across the junctions and the Hi-C contact maps, ruling out potential mis-assembly for this chimeric chromosome (**Fig. S4**). Moreover, there were major genomic inversion events inside Chr06 and at each end of Chr07 in FonR2 compared to the Fo47 reference (**Fig. 2B**), whereas FonR1 and FonR3 only had some minor inversion events.

### Comparative genomics reveals race-unique accessory chromosomes/regions

Given the distinctive virulence profiles of Fon races on differential hosts, their genomes likely harbor unique DNA elements that determine their race-specific virulence traits. To identify such elements, we performed a synteny analysis of the three genomes and found that they shared a high level of synteny (82.78% synteny on average whole genome) in both core chromosomes and accessory chromosomes (**Fig. 3**). In addition to the 11 conserved core chromosomes (average 84.77% gene synteny among three races’ CC), some accessory chromosomes were also syntenic between three races. For example, synteny was observed for accessory chromosome Chr07 of FonR1 and FonR3, and the accessory regions on Chr04 in FonR2. GO enrichment analysis showed that transcription regulation-related genes were enriched in these chromosomes and regions in all isolates (**Fig. S6**). To compare the gene compositions of the three genomes, we performed gene homology analysis on the predicted gene models using OrthoFinder^29^. We identified 16,464 orthogroups across three races, among which 11,513 were single-copy orthogroups. There were 391, 1113, and 464 race-specific genes in FonR1, FonR2, and FonR3, respectively. 59.1% (231/391), 31.6% (352/1113), and 48.9% (227/464) of race-specific genes in FonR1, FonR2, and FonR3 were found in race-specific accessory regions (**Fig. S7**).

**Fig. 3.**
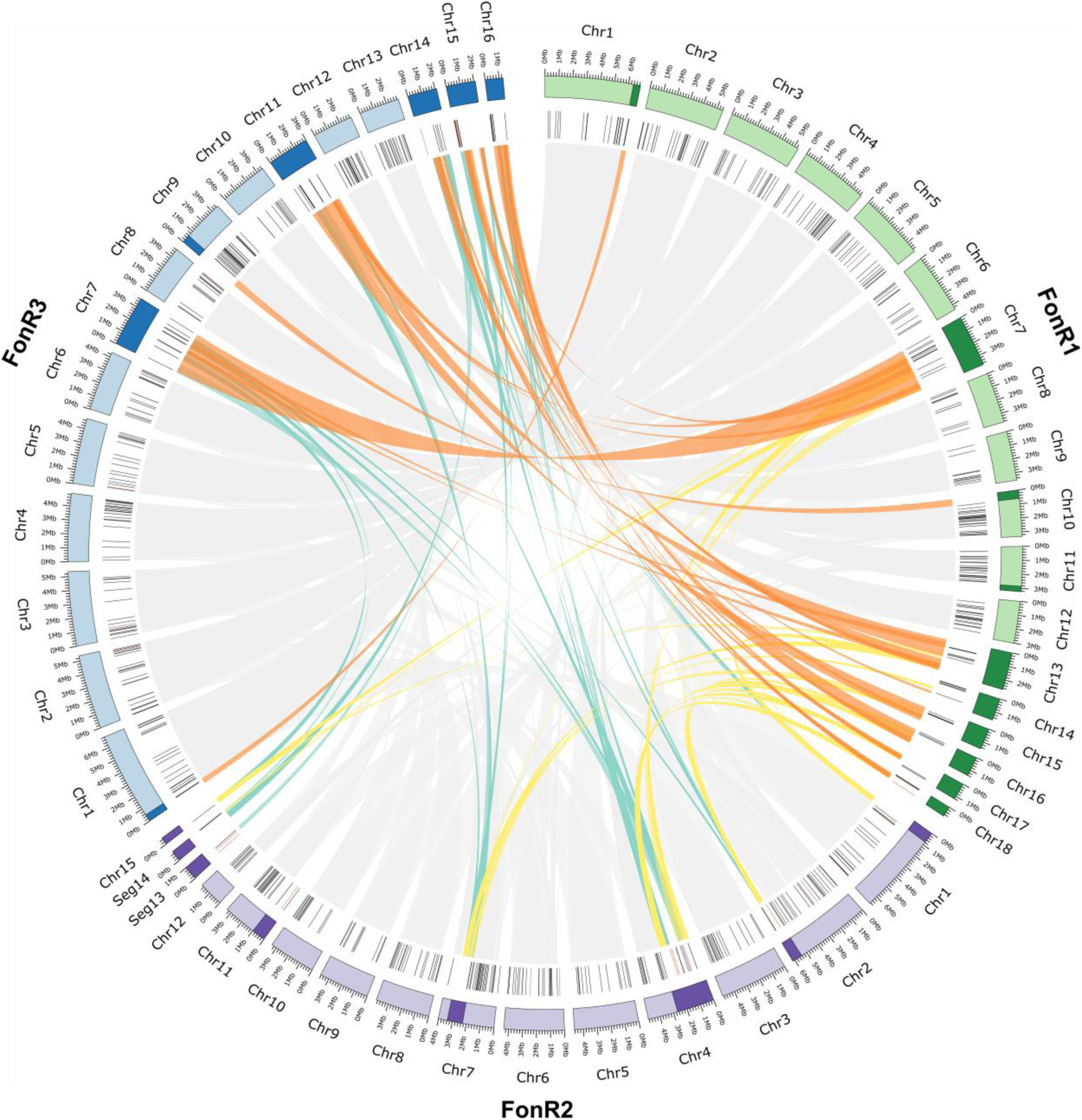
Comparative genomics of FonR1, FonR2, and FonR3 gap-free genome assemblies reveals core and lineage-specific accessory regions/chromosomes. (i) Chromosomes of FonR1, FonR2, FonR3. Light colors represent core chromosomes (CC), while dark colors represent accessory chromosomes/regions (AC/AR). (ii) Predicted effectors (black) and unique effectors (red) are shown. (iii) Links showing synteny between three genome sequences. Gray links are between CCs while colored links are between AC/ARs.

As accessory genomes are hotspots of virulence genes likely contributing to race evolution in *F. oxysporum*, we investigated the gene expression of core and accessory genomes in three Fon races during the infection of watermelon plants susceptible cultivar G42. While the global gene expression shows little skewness in response to Fon infection (median log2FC = 0.137-0.195), the expressions of accessory genes and putative effectors were skewed towards up-regulation by infection (median log2FC = 0.891-1.406), suggesting more induced expression of accessory genes) (**Fig. 4A**). We then identified differentially expressed genes (DEGs) between the control and 6 dpi samples. We observed significantly more infection-stage DEGs from accessory genomic regions than from core ones. For example, 15.36% (1217/7922), 15.19% (1245/8194), and 14.10% (1166/8267) of the genes in core genomes were significantly up-regulated during infection in FonR1, FonR2, and FonR3, respectively. By contrast, 27.27% (177/649), 24.14% (127/526), and 25.50% (166/651) of the genes in accessory genomes were significantly up-regulated during infection (**Fig. 4A**). These observations indicated that a higher portion of genes in accessory regions were up-regulated during infection than the core regions, confirming their roles in Fon infection. Among the single-copy orthologues, 10,006 genes were expressed in at least one condition and only 1944 (16.9%) genes exhibited different expression patterns among the three races (**Fig. S7**), suggesting that the expression of the orthologous genes was mostly similar, despite showing some divergence.

**Fig. 4.**
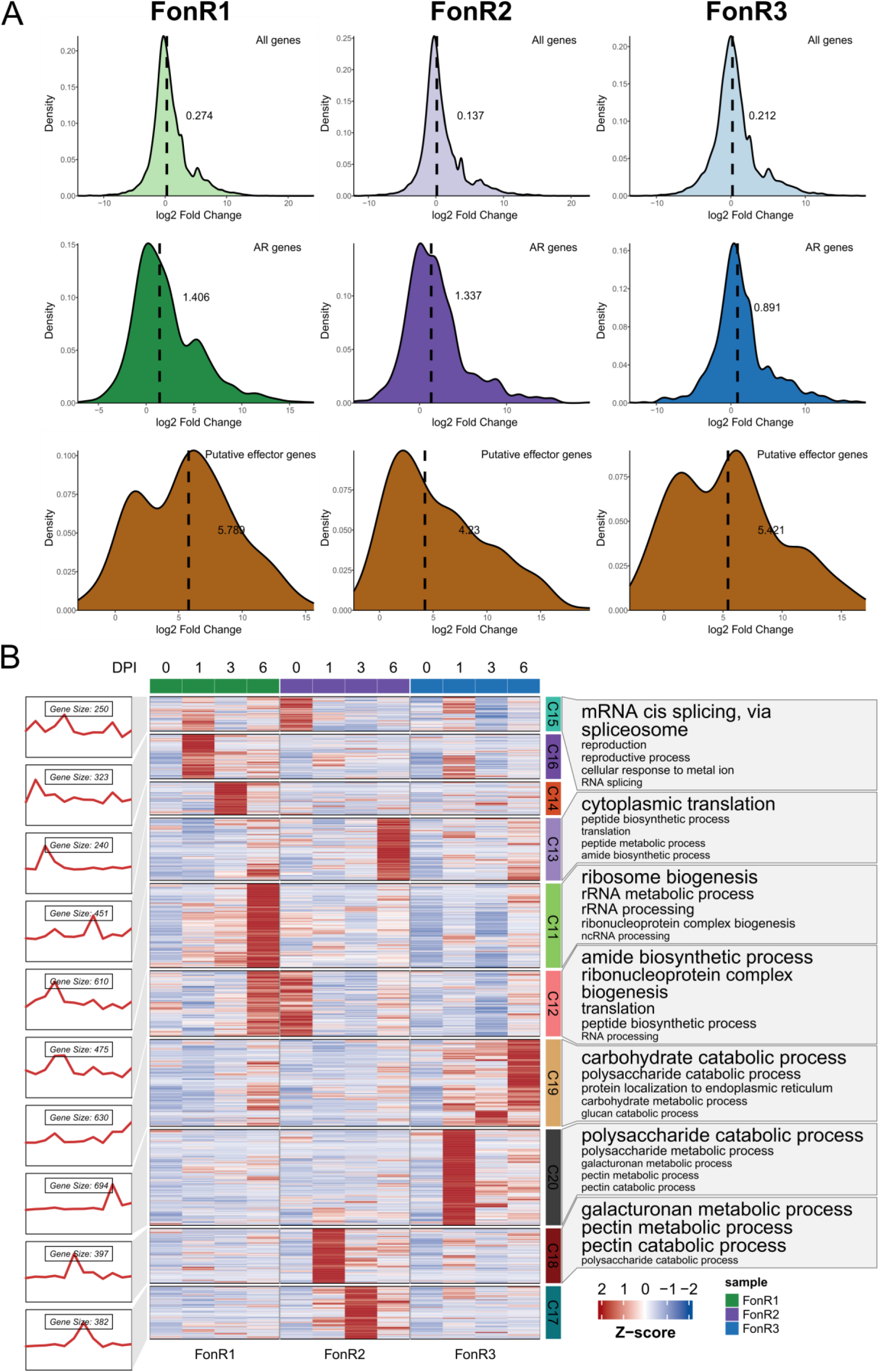
Comparative transcriptomics of three Fon races during watermelon infection. (A) Kernel density plots of gene expression log2-fold change values between control and 6 days post-inoculation of the G42 cultivar. The median log2-fold change values are shown as bold numbers and dashed vertical lines. (B) k-means clustering of the expression of the single-copy orthologous genes of FonR1, FonR2, and FonR3. The figure was generated using ClusterGVis^70^ with k=20. Only the clusters with a difference among the three isolates are shown. The full image, including all clusters, can be found in **Fig. S5**.

### Transcriptome clustering reveals race-conserved and divergent gene expression modules

To detect the differences in expression patterns in the conserved genes shared by FonR1, FonR2, and FonR3, we performed k-means clustering (k=20) on the TPM expression matrix of the expressed single-copy orthologs (SCOs) (**Fig. 4B; Fig. S8**). According to the results, the genes in Cluster 20 (C20) and C19 are up-regulated only in the FonR3 post-inoculation samples. Particularly, SCOs in C20 exhibited expression patterns of earlier and higher up-regulation in planta in FonR3 and were enriched with GO terms including "polysaccharide metabolic process", "galacturonan metabolic process", and "pectin metabolic process", suggesting their involvement in plant cell wall degradation. In addition, C20 contained an SCO encoding a pectate lyase C "FonR3Chr10G008700" reported as a plant virulence factor previously^30^. C19 with SCOs enriched in "carbohydrate catabolic process" showed higher upregulation in FonR3 than FonR1 and FonR2. Significantly, three SCOs "FonR3Chr12G002170", "FonR3Chr09G004770", and "FonR3Chr01G010130" encoding glycosyl hydrolase 5, 6, and 7, respectively, exhibited a 768 to 1120-fold expression increase in FonR3 than in FonR1 and FonR2 (65 to 177-fold change) during infection (**Fig. 4B**). The different expression pattern of SCOs across three races indicated that FonR3 might be equipped with different transcription regulators than FonR1 and FonR2 during infection. Furthermore, to study the expression patterns of genes unique in each race, we performed separate k-means clustering (k=20) for FonR1, FonR2, and FonR3 using their gene expression matrices, followed by functional enrichment analysis. Interestingly, the GO term "polysaccharide metabolic process" indicative of plant cell degradation was enriched in clusters of all three races but was present in clusters with different expression patterns (C18 in FonR1, C3 in FonR2, and C17 in FonR3 in **Fig. S9-11**). For example, for FonR1 and FonR2, genes in the "polysaccharide metabolic process” enriched clusters were up-regulated at 3 and 6 dpi. However, in FonR3, this up-regulation began at an earlier stage of infection (1 dpi). This indicated that the polysaccharide metabolic process regulation might play a part during infection and the differences in the gene expression patterns of this process might be associated with the differences in virulence.

### Effectorome analysis reveals putative race-specific effectors (RSEs) in FonR3

Effectors, defined as small secreted proteins, are crucial to the successful infection of plant hosts for *F. oxysporum*^31,32^. Considering the importance of effectors, we comprehensively predicted the putative fungal effectors in three Fon races using a pipeline involving three tools commonly used in fungal secretome and effector identification: EffectorP^33^, SignalP^34^, and TMHMM^35^. The candidate effectors were identified by overlapping EffectorP and SignalP results and removing TMHMM hits, resulting in 467 to 489 candidate effectors in three races (**Fig. S5**). Furthermore, we screened out the putative effectors that were highly expressed in post-inoculation samples and little or no expression in control samples (fungal cultures) based on expression patterns derived from hierarchical clustering of transcriptome data, identifying 255, 247, and 263 putative effectors in FonR1, FonR2, and FonR3, respectively, located both on core and accessory chromosomes (**Fig. 2C**). Captured in the putative effectors of three Fon races was *FonSIX6* (FonR1Chr18G000080, FonR2Chr08G000200, FonR3Chr15G000080), the Fon homolog of the well-known avirulence effector *SIX6* (secreted in xylem protein 6)^36^. K-means clustering of SCOs suggested these putative effectors are distinctively regulated in three races during watermelon infection, where 67 of 240 single-copy effectors belonging to two clusters (C19 and C20) were only upregulated in

#### FonR3 during early infection

So far, no resistant watermelon germplasm resources or commercial cultivars to Fon race 3 have been reported^8^, therefore Fon race 3 poses a potential and serious threat to future watermelon production worldwide despite its sporadic reports. Given the importance of Fon race 3, our study focused on identifying and dissecting the race-specific virulence factors, particularly functional effectors that enabled Fon race 3 to overcome the resistance in cultivar PI296341 and PKR6. Comparative genomic analyses showed that 11, 10, and 13 putative effectors were unique to FonR1, FonR2, and FonR3, respectively (**Fig. S5**). We conducted an in-depth analysis of the 13 race-specific effectors (RSEs) that were only present in FonR3 compared to FonR1 and FonR2. It is noteworthy that the majority (10) of the 13 FonR3 RSEs are situated on core chromosomes, with three located in accessory regions (**Fig. 3**). Transcriptome data showed that all 13 RSE genes were expressed at much lower levels during the vegetative growth stage but highly expressed *in planta* (**Table S1**). Moreover, most mature proteins of 13 RSEs had fewer than 224 amino acid residues, rich in cysteines, and possessed N-terminal signal peptides (**Table S1**).

### FonRSE1 is a critical effector for FonR3 virulence

To study the functions of putative RSEs of FonR3, we generated the gene deletion mutants via the gene replacement approach in the wild-type (WT) FonR3 and obtained 8 of 13 RSE mutants (**Table S1-2**). In a preliminary greenhouse bioassay on watermelon cultivars, seven of eight candidate RSEs’ mutants showed varied degrees of reduced virulence compared with WT FonR3, among which △*FonRSE1* was the least virulent, leading to weakened disease symptoms on PKR6 seedlings, and postponed plant death for Sugarlee and G42 seedlings (**Fig. S12**). With the gene located in core chromosome Chr2 (FonR3Chr02G017230), FonRSE1 is a very small protein with 67 amino acids consisting of a signal peptide and an unknown domain, aligned with the non-conservative nature of functional domains of effectors^37^ (**Fig. 5A**). The transcription of *FonRSE1* maintained at a fairly low level during vegetative growth (0 dpi) but was significantly upregulated after infection initiated (1, 3, 6 dpi) (**Fig. 5B**; **Table 1**). The predicted 3D structure of FonRSE1 by AlphaFold2-based ColabFold open-source software^38^ had an *α*-helix, three *β*-sheets, and several flexible-loop regions (**Fig. 5C**). Cysteine residues of fungal effectors could form multiple disulfide bonds to promote the proper protein folding and stability^39^. FonRSE1 carries three cysteine residues, which were predicted to form one disulfide bond between cysteine 42 and cysteine 63 (**Fig. 5C**).

**Fig. 5.**
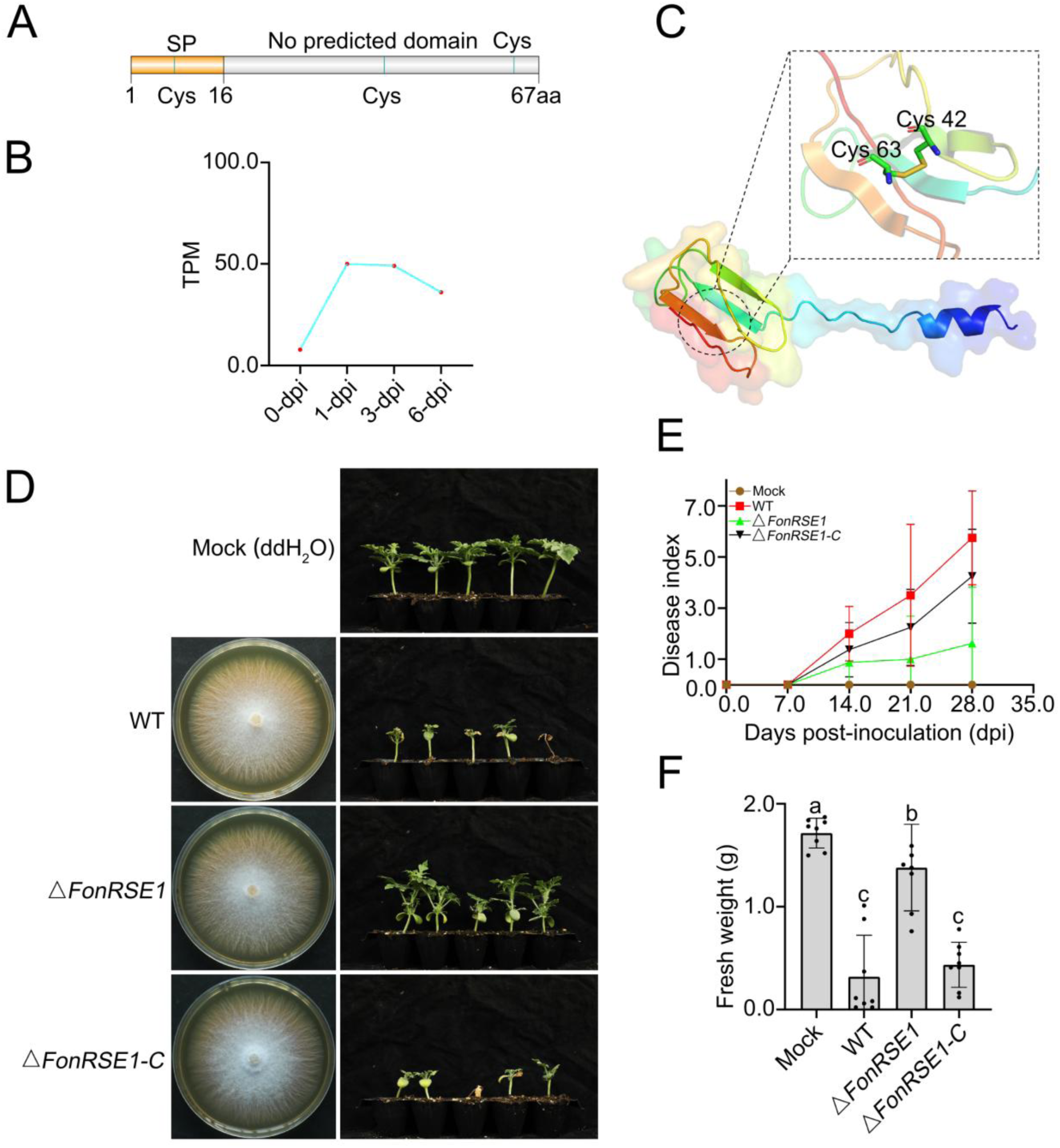
Features of FonRSE1, a critical effector for FonR3 infection. (A) A structure diagram for FonRSE1 with domains and cysteine residues. (B) The expression of *FonRSE1* in the vegetative growth stage (0 dpi) and in planta (1, 3, 6 dpi). (C) Predicted 3D protein structure of FonRSE1 using Alphafold2 and ColabFold software^38,78^. (D) Greenhouse bioassays for the functional study of *FonRSE1* were repeated independently three times, with eight to ten plants for each treatment. One representative experiment was shown here. Left lane: colony growth of WT FonR3, the *FonRSE1* mutant, and its complemented strain on potato dextrose agar (PDA) plates at 5 dpi; Right lane: Infected PKR6 plants with mock (water) and the corresponding Fon strains from the left lane at 28 dpi using 11-day-old seedlings. (E) Disease index quantification of infected PKR6 seedlings from 7 to 28 dpi. Disease index was evaluated based on a 7-scale rating: 0 = asymptomatic, 1 = slight stunted growth and yellowing, 3 = stunted growth and yellowing, 5 = wilting, 7 = dead. (F) Fresh weight measurements of the above-ground infected plants at 28 dpi. Different letters above the bars represented the significant difference between treatments using ANOVA analysis followed by Duncan’s multiple range test (*p* = 0.05).

To further validate the function of *FonRSE1,* we generated a complemented strain △*FonRSE1*-*C* for △*FonRSE1*. The mutant and complemented strain generated showed no significant differences with WT FonR3 on growth rates, colony morphology, and conidiation, implicating that the absence of *FonRSE1* had no detectable interference with the fungal vegetative growth (**Fig. 5D; Fig. S13**). The greenhouse bioassays using △*FonRSE1* and △*FonRSE1*-*C* were repeated three times, with eight to ten plants per treatment (**Fig. 5D; Table S3; Fig. S14**). Besides occasional quick death around 7 dpi, the majority of inoculated PKR6 plants by WT FonR3 started to show leaf yellowing and stunted growth after 7 dpi, and the symptoms continued to develop for 4 to 8 weeks post-inoculation until complete plant death. In one representative experiment, infected PKR6 plants by WT FonR3 weighed 0.32 g while mock plants weighed 1.72 g on average at 28 dpi. (**Fig. 5E-F**). The deletion of *FonRSE1* led to a strong alleviation of the disease index with a significantly increased average fresh weight of 1.38 g of PKR6 seedlings at 28 dpi. The complemented strain △*FonRSE1-C* provoked a comparable level of disease index to WT and an average fresh weight of 0.44 g, indicating a restoration of virulence. Therefore, we inferred that FonRSE1 was an important virulence effector for the FonR3 to overcome the previously resistant cultivar PKR6. Though unique to FonR3 and absent in FonR1 and FonR2 isolates, *FonRSE1* shares 100% identity with a hypothetical protein (accession KAF6523144.1) of *F. oxysporum* f. sp. *conglutinans* (Foc) strain Fo5176^40^, and 97% identity with an uncharacterized protein (KAJ0153318.1) of *F. oxysporum* f. sp. *albedinis* (Foa). Taken together, it indicated that a reservoir for FonRSE1 effector in FOSC might be the source for the emergence of Fon race 3.

## DISCUSSION

The asexual filamentous fungus *F. oxysporum* is the causal agent of vascular wilt in over 100 different crop species, a major threat to agricultural productivity with few effective control measures. Genomics is the key to understanding the mechanisms behind the pathobiology, especially the rapid evolution of pathogen virulence. Despite their small genomes, it is challenging to fully assemble their genomes due to repeat-rich chromosomes. The genomes of multiple members in FOSC have been reported since the first release of the tomato pathogen *F. oxysporum* f. sp. *lycopersici* (Fol4287)^3^, revealing their characteristic two-speed genome structures with horizontally transferred accessory chromosomes. Little is known about the host-pathogen mechanisms of Fon, the causal agent of watermelon vascular wilt of watermelon, due to the lack of high-quality chromosome-level reference genomes. While the genome of different races of Fon was assembled previously^18^, these releases are contig-level and fragmented, unable to reveal the chromosome structure and evolution via comparative genomics. Here, we combined the cutting-edge long-read (PacBio and ONT) sequencing and Hi-C sequencing to assemble and annotate the gap-free genomes for three isolates of Fon, representing three physiological races of Fon showing distinctive virulence towards different hosts. All assembled chromosomes were gapless with all centromeres and most telomeres captured, except for FonR2, which missed 17 telomeres. Before this study, only two gap-free genome assemblies were released for *F. oxysporum*, the endophyte Fo47^28^ and *Arabidopsis* pathogen Fo5176^41^. Therefore, we expect that the three gap-free Fon genomes from this study will be a great addition to the high-quality genome resources of FOSC to study their pathobiology and genome evolution.

Genome variation is a major driving force for evolution and the generation of new traits. Rapid evolution is a hallmark of fungal pathogens, especially *F. oxysporum,* allowing them to adapt to new environments and resistant cultivars. The different races, especially race 3, of Fon have overcome all known Fusarium wilt-resistant watermelon germlines. Our analysis of the three gap-free genomes yielded a couple of surprising findings. First, we are surprised by the observation of a large difference in their genome structure, such as the number of chromosomes, even at the race level within the same *F. oxysporum* formae speciales. Second, it is also surprising to see the higher sequence similarity and collinearity between race 1 and race 3 than each with race 2, highlighting the uncanonical and complex evolutionary trajectories of the Fon genome and race evolution. Such karyotype difference and the polyphyletic nature of Fon races hinted potential independent origin and evolution of fungal virulence on watermelon hosts, likely involving a complex history of genetic introgression and recombination. By comparing the gap-free genomes, we dissected the conserved and distinctive genome characteristics behind race evolution, and identified the accessory chromosomes and regions in all three races, which showed substantial synteny with high sequence identity, but also had unique accessory genomic regions. Comparison of annotated genes revealed largely conserved gene families with similar expression patterns, reflecting the core gene regulatory machinery for Fon. However, divergence of expression was also observed for conserved genes with and without interaction with host plants, suggesting an evolution of cis- or trans-regulatory elements for these highly conserved coding regions. For example, K-means clustering of single-copy ortholog expression profiles revealed gene modules (C19 and C20) up-regulated exclusively in FonR3 post-inoculation (**Fig. 6**). The fact that most of these genes located on core genomic regions also indicated that genome and transcriptome variation in core regions also likely contribute to distinctive virulence of races on hosts, although experiments are needed to study these orthologs with diverged expression in future studies of pathogen evolution which can focus on mapping the regulatory elements and identifying functional genomic variants using population genomic data.

**Fig. 6.**
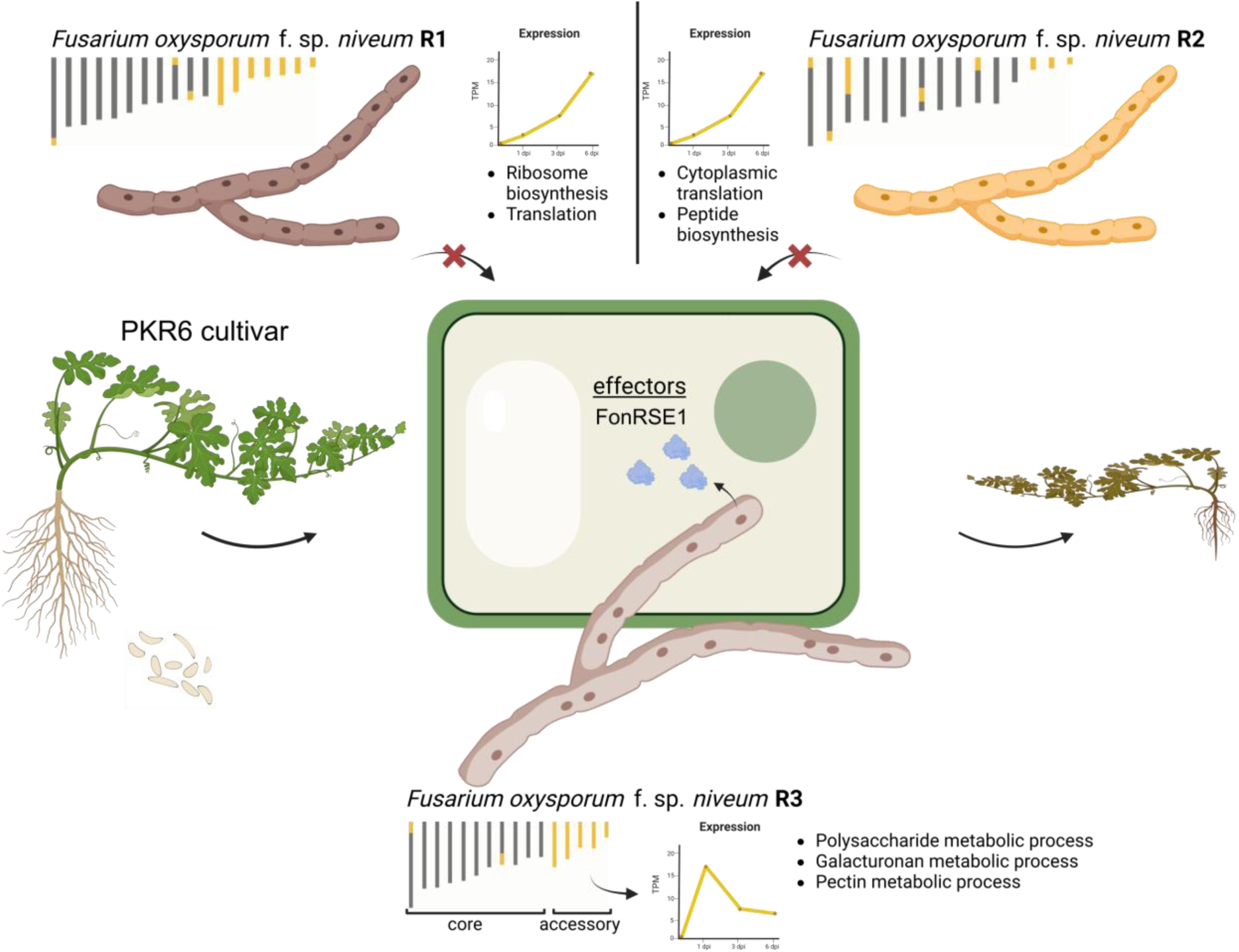
A schematic diagram summarizing the evolution of virulence in *F. oxysporum* f. sp. *niveum* (Fon). Core (gray) and accessory (yellow) chromosomes/regions of FonR1, FonR2, and FonR3, and the accessory genome expression patterns with the enriched GO terms in infection on the susceptible G42 cultivar are shown. While FonR1 and FonR2 cannot infect the resistant cultivar PKR6, FonR3 can infect it and cause wilt symptoms through the RSE1 effector. This image was created by Biorender.

Theoretically, vital effectors determining host specificity should be largely shared across all Fon isolates. The putative effector profiles of FonR1 (255), FonR2 (247), and FonR3 (263) predicted in this study provided valuable resources for further functional study of Fon effectors. As more high-quality genome assemblies of Fon races from different geographic locations and their putative effector profiles become available, the vital effectors determining host specificity can be further narrowed down and tested through gene editing or mutagenesis. The identification of 13 RSEs in FonR3 highlights critical factors contributing to its pathogenicity on a broad spectrum of watermelon differential cultivars. The mutagenesis experiment of *FonRSE1* confirmed its critical role for FonR3 to overcome previously resistant cultivar PKR6 (**Fig. 5**; **Fig. 6**). Though absent in FonR1 and FonR2, *FonRSE1* is conserved in other ff. spp. such as Foc and Foa. Given the polyphyletic nature of Fon and FOSC in general, and the fact that strains from different ff. spp. could be found genetically closer than strains of the same f. sp.^3,16,42^, it is not surprising that effectors can be shared between ff. spp. Therefore, new pathogenic races may arise through acquiring new effectors vertically and horizontally from effector reservoirs throughout the FOSC.

As the genome composition of FOSC is highly diverse and polyphyletic, it is likely that each Fon race, identified using host differentials, might represent a population of fungal strains with similar genetic makeup. Isolates belonging to the same race but from different geographic regions may differ greatly in their genome compositions, such as chromosome numbers or effector repertoire. As a result, the limitation of this study is that one gap-free genome assembly may not capture all genetic variations of its designated race. A thorough understanding of Fon races would be further advanced by the availability of Fon population or pan-genomics using high-quality genomes in the future.

In summary, the three gap-free genome assemblies, annotations, and the RSEs of Fon physiological races are valuable genetic resources for dissecting *F. oxysporum* pathobiology on watermelon, studying fungal genome evolution, and improving development of effective disease control strategies.

## Materials and Methods

### Fungal isolates and plant materials

Three Fon races (FonR1, FonR2, FonR3) and seeds of watermelon cultivars (G42, Calhoun Gray, Sugarlee, and PKR6) were kindly provided by the Crop Genetics and Breeding Platform of Peking University Institute of Advanced Agricultural Sciences, Weifang, Shandong. FonR3 (NXWFW008) was recently isolated from infected watermelon plants in Ningxia Province, China^9^. Fungal isolates, including derived mutants generated in this study (**Table S1**), were routinely cultured on Potato Dextrose Agar (PDA) plates at 26°C. Conidiation was measured with 5-day-old Potato Dextrose Broth (PDB) cultures, and conidial germination was assayed with liquid Yeast extract peptone dextrose (YEPD) broth cultures (0.3% yeast extract, 1% peptone, and 2% dextrose) at 26°C in a rotary shaker at 175 rpm for 12 h^43^. For ultra-long genomic DNA isolation, the vegetative hyphae were harvested from 2-day-old PDB cultures set at 175 rpm and extracted using a cetyltrimethylammonium bromide (CTAB) method^44^.

### Greenhouse bioassays

Seeds of watermelon cultivars were surface sterilized in 0.1% mancozeb for 30 minutes before rinsing with distilled water. To improve the germination rate, disinfected seeds were incubated at 30°C for 48 h^10^. Then germinated seeds were sown in perlite and placed in a growth room set at 26°C/16h and 18°C/8h with a humidity of 80%. After 11 days, seedlings were uprooted and washed. Roots were dipped into spore suspensions of 5×10^6^ spores/ml for one minute^10,43^ and the inoculated seedlings were planted in sterilized soil (substrate/perlite/vermiculite = 3:1:1) and maintained in the above-growth room for four weeks. Disease phenotypes were monitored and documented once a week according to a disease index scale: 0 = asymptomatic, 1 = slight stunted growth and yellowing, 3 = stunted growth and yellowing, 5 = wilting, and 7 = dead. Plant tissues above the soil were harvested at four weeks post-inoculation and fresh weight was measured immediately after harvesting.

### Genome and transcriptome sequencing

Illumina paired-end DNA sequencing library was prepared using NEB Next® Ultra™ DNA Library Prep Kit for Illumina (NEW ENGLAND BioLabs) following the manufacturer’s instructions. The 150 bp paired-end reads were produced using the Illumina Novoseq 6000 platform by Novogene Biotechnologies Inc. (Tianjin, China). PacBio SMRTbell library was constructed using PacBio SMRTbell Express Template Prep Kit 2.0 (PacBio). Short reads were removed with a 15 kb cutoff on BluePippin (Sage Science). HiFi consensus reads were generated using a PacBio Sequel II system at Novogene Biotechnologies Inc. (Tianjin, China). The Nanopore DNA library was prepared following the Ligation Sequencing SQK-LSK109 Kit (Oxford Nanopore Technologies, Oxford, UK) protocol and sequenced using the Oxford Nanopore GridION (20 kb) platform. The Hi-C library was prepared from cross-linked chromatins with a standard Hi-C protocol^45^. Then the library was sequenced using Illumina NovaSeq 6000 to obtain 150 bp paired-end reads at Novogene Biotechnologies Inc. (Tianjin, China). Total RNA was extracted using Trizol Reagent (Thermo Fisher Scientific). The mRNA was subjected to transcriptome sequencing library construction using the Illumina TruSeq transcriptome kit (Illumina). The libraries were then sequenced using the Illumina Novaseq 6000 platform at Biomarker Technologies Corporation (Qingdao, China) to generate 150 bp paired-end reads.

### Genome assembly

Genome sizes of the three isolates were estimated using the Illumina data by Jellyfish v2.3.0^46^ (k-mer size = 19) and Genomescope v1.0^47^. HiFi and ONT reads were assembled with Hifiasm v0.19.5^21^. To remove within-species contamination and acquire the nuclear genome, we aligned the contigs of the initial assembly to the reference genome of *Fusarium oxysporum* mitochondria (GenBank accession: NC_017930) with Minimap2 v2.24^48^. Contigs with above 50% coverage were removed. To remove bacterial contamination, we ran a megaBLAST^49^ search against a database of common contaminants in eukaryotic genome assemblies (ftp://ftp.ncbi.nlm.nih.gov/pub/kitts/contam_in_euks.fa.gz) and the reference genome of the contamination bacteria (*Brucella intermedia*, GenBank accession: GCA_900454225.1) we detected in FonR2 samples. Then we remove the contamination contigs with the following criteria^50^: requiring e-value ≤ 1×10^−4^, reporting matches with ≥ 98% sequence identity and match length 50–99 bp, ≥ 94% and match length 100–199 bp, or ≥ 90% and match length 200 bp or above. To anchor the contigs to chromosomes, we applied the pipeline of Juicer v1.5^22^, 3D-DNA v180419^51^, and Juicebox v1.11.08^23^. The anchored scaffolds were manually examined and adjusted within Juicebox for assembly validation. To assess genome completeness, we applied BUSCO v5.4.3^25^ for ortholog detection using fungi_odb10 database. Quality values (QV) based on HiFi reads were estimated using Merqury v1.3^52^ with the k value of 23.

### Genome annotation

To annotate repeats in the genome assemblies, we used the universal Repbase database and a species-specific de novo repeat library constructed by RepeatModeler v2.0.6^53^ to annotate the DNA sequences and then annotated and masked the repetitive elements by RepeatMasker v4.1.2^27^. To predict gene models, we assembled the RNA-Seq reads with HISAT2 v2.1.0^54^ and Stringtie v2.2.1^55^ for transcriptome evidence and downloaded the protein sequences of *Fusarium oxysporum* and the universal fungi protein knowledgebase from Swiss-Prot^56^ for homologous protein evidence. Transcriptome-based prediction, protein homology-based prediction, and *ab initio* prediction were combined in the pipeline of MAKER v3.01.03^26^ and BRAKER v2.1.6^57^. We first trained the GeneMark-ET and AUGUSTUS models using BRAKER and trained the SNAP semi-HMM model using MAKER. Then we integrated the trained SNAP, GeneMark-ET, and AUGUSTUS models into MAKER to predict more credible genes. Finally, the highest-ranking gene sets were retained based on Annotation Edit Distance (AED < 0.5)^58^. Gene functional annotation was conducted using eggNOG-mapper v2.1.12^59^.

### Fungal effector prediction

To select the putative fungal pathogenic effector proteins from the predicted gene models, we first used the neural network of SignalP v6.0^34^. The predicted signal peptides were examined with TMHMM v2.0^35^ and the peptides with detected transmembrane helices were excluded from the predicted secretome dataset. We then predicted the effector proteins from the putative secretomes using EffectorP v3.0^33^. To further screen the putative effectors, quantifications of gene transcripts from RNA-seq reads of axenic and post-inoculation samples were performed using kallisto v0.48.0^60^. The putative effectors with higher expression levels in post-infection samples were selected using the hierarchical clustering in R base functions for experimental verification.

### Gene homology and synteny analysis

Orthologues and orthogroups were inferred using OrthoFinder v2.5.4^29^ with default value settings and ‘-M msa’ activated. We identified the genes in race-specific orthogroups as race-specific or race-unique genes. The phylogenetic tree was generated using the species tree by OrthoFinder using 18 publicly available *Fusarium* genomes and drawn using iTOL v5^61^. The gene synteny analysis was performed by JCVI v1.1.19^62^. The syntenic gene blocks were identified by performing an all-against-all LAST search and chaining the hits with a distance cutoff of 20 genes and a minimum block size of 5 gene pairs.

### Transcriptome analysis

Raw transcriptome data were pre-processed with fastp v0.23.2^63^ to trim artifacts and improve data quality. Gene expression levels were then quantified using kallisto v0.48.0^60^. Counts for mapped reads were normalized by TPM (transcripts per million). To visualize the expression patterns of the focused genes among samples, heatmaps were generated using the R package ComplexHeatmap v2.12.1^64^. Gene function and protein homology term enrichment analyses based on gene ontology (GO)^65,66^ and PFAM^67^ databases were performed using ClusterProfiler v4.8.3^68^. The differential expression analysis was performed using the R package DESeq2 v1.40.2^69^. We identified genes with log2 fold changes greater than or equal to 2 and adjusted p-values less than 0.05 as significantly up-regulated genes, and genes with log2 fold changes less than or equal to -2 and adjusted p-values less than 0.05 as significantly down-regulated genes. The k-means clustering of gene expression patterns was conducted and visualized using ClusterGVis v0.1.1^70^.

### Generation of knock-out mutants for FonR3 RSEs

Protoplast preparation and polyethylene glycol (PEG)-mediated transformation were performed as described previously^71^. Briefly, hyphae of FonR3 grown in YEPD for 12 h were lysed with an enzyme lysis buffer containing driselase (250 mg/ml; Sigma-Aldrich, St. Louis, MO, USA) and lysing enzymes (50 mg/ml; Sigma-Aldrich, St. Louis, MO, USA) under a condition of 31°C, 110 rpm for 2-3 h. Then released protoplasts were collected by centrifugation at 3,500 rpm at 4°C^43^. Protoplasts were subsequently washed twice with STC buffer (20% sucrose, 50 mM CaCl_2_, 10 mM Tris pH 7.5) and resuspended to a final concentration of 10^5^ - 10^6^ per ml with STC buffer.

Gene replacement constructs of specific effectors in the FonR3 isolate were generated using a split-marker approach^72^. or each effector gene, approximately 1.0 kb 5’ upstream and 3’ downstream regions were respectively amplified from FonR3 with gene-specific primer pairs 1F/2R and 3F/4R (**Table S2**). The resulting PCR products were ligated to the 5’- and 3’-regions of the hygromycin phosphotransferase (*hph*) cassette by overlapping PCR^73,74^.

For transformation, 200 μl protoplasts and 5 μg fused constructs were mixed and incubated at 26°C for 20 min, followed by supplementing 1 ml PSTC (40% PEG-8000 in STC buffer) and another 20-min incubation. Then 10 ml regeneration medium (0.3% yeast extract, 20% sucrose, 0.3% casein enzymatic hydrolysate) was added to the mixture before an overnight incubation in a shaker set at 110 rpm and 26°C. The resulting transformants were selected on Top agar medium (0.3% yeast extract, 0.3% casamino acids, 20% sucrose, and 1.5% agar) with 300 μg/ml hygromycin B (CalBiochem, La Jolla, CA, USA). Positive transformants were identified by PCR using corresponding primers (**Table S2**).

### Generation of the complemented strain

For complementation of the deletion mutant △*FonRSE1*, a strong promoter from FonR3 (promoter of FonR3Chr10G008340), the *FonRSE1* gene with its 1.5 kb downstream region was amplified with its specific primers (**Table S2**) and ligated by overlapped PCR. The ligated fragments were cloned into XhoI-digested pFL2 vector by the yeast gap repair approach using yeast strain XK1-25^75,76^. The recombined constructs carrying the geneticin-resistant marker were rescued from Trp^+^ transformants and transformed into the protoplasts of the mutant. Transformants containing the complementation constructs were screened with geneticin and identified by PCR assays.

## ACKNOWLEDGMENT

We would like to thank the Bioinformatics Platform at Peking University Institute of Advanced Agricultural Sciences for providing high-performance computing services. We would also like to thank Dr. Li-Jun Ma from the University of Massachusetts Amherst for her valuable discussions.

## AUTHOR CONTRIBUTIONS

L.G. conceived and supervised this project. Y.D. and X.Z. provided the fungal isolates and plant seeds for this study. H.W. and L.Z. prepared materials for sequencing. D.H.A., S.Y., and X.R.W conducted genome assembly. D.H.A. and S.Y. performed genome annotations, comparative genomics and transcriptomics analysis. X.W handled genomic data curation and submission, and participated in discussion. H.W., L.Z., G.W., D.M., L.X., X.G., L.W., and Q.Y. performed experiments. D.H.A., L.G., and L.Z. prepared the main figures and tables, wrote and revised the manuscript. H.W., S.Y., and Z.K. assisted in the preparation of figures. All authors read and approved the manuscript.

## FIGURE LEGENDS

**Fig. S1. Hi-C contact maps of FonR1, FonR2, and FonR3 genome assemblies.**

**Fig. S2. Confirmation of gap closing in FonR2 assembly.** On the left the chromosomes with gaps shown in black squares in the Hi-C contact maps, and on the right, IGV screenshots of HFi and ONT read mappings at the gap region.

**Fig. S3. Genome alignment of FonR1, FonR2 and Fo47.** Ideograms of FonR1 (A) and FonR2 (B) were visualized using GenomeSyn^77^. Genomic features, including gene density, GC content, TE content, telomeres, centromeres, and genome synteny with Fo47 chromosomes are shown on the ideograms. The yellow background indicates the accessory regions.

**Fig. S4. Confirmation of AR/CC border in FonR2.** Hi-C contact map showing Chr07 of FonR2 on the left and IGV screenshots with HiFi and ONT reads mappings on the AR/CC borders. AR: accessory regions. CC: core chromosome

**Fig. S5. Fon effector prediction pipeline.** The effectors are first identified if they are predicted by SignalP^34^ and EffectorP^33^ but not including the transmembrane domain predicted by TMHMM^35^. After filtering, only the expressed genes in any of the selected conditions for transcriptome analysis, only genes that have increased expression *in planta* have been selected as putative effectors. Finally, race-specific effectors were identified with OrthoFinder^29^.

**Fig. S6. Gene ontology (GO) enrichment analysis of accessory chromosomes (AC) of FonR1 (left), FonR2 (middle), and FonR3 (right).** The color of bubbles is scaled by the adjusted p-values from enrichment analysis. The size of bubbles is scaled by the number of genes annotated with the specific GO terms. BP: biological process. CC: cellular component. MF: molecular function

**Fig. S7. Donut plot summarizing the ratio of the ortholog gene family composition and gene expression conservation in annotated genes of three Fon races.**

**Fig. S8. Heatmap showing the transcriptome clustering using K-means (k=20) clustering of single-copy ortholog expression of three races (FonR1, FonR2, and FonR3) at fungal culture and during infection of watermelon cultivar G42 at different time points (days) after inoculation (D1AI, D3AI, and D6AI).** Significantly enriched gene ontology terms are listed on the right side.

**Fig. S9. Heatmap showing the transcriptome clustering using K-means (k=20) clustering of FonR1 expression at fungal culture and during infection of watermelon cultivar G42 at different time points (days) after inoculation (D1AI, D3AI, and D6AI).** Significantly enriched gene ontology terms are listed on the right side.

**Fig. S10. Heatmap showing the transcriptome clustering using K-means (k=20) clustering of FonR2 expression at fungal culture and during infection of watermelon cultivar G42 at different time points (days) after inoculation (D1AI, D3AI, and D6AI).** Significantly enriched gene ontology terms are listed on the right side.

**Fig. S11. Heatmap showing the transcriptome clustering using K-means (k=20) clustering of FonR3 expression at fungal culture and during infection of watermelon cultivar G42 at different time points (days) after inoculation (D1AI, D3AI, and D6AI).** Significantly enriched gene ontology terms are listed on the right side.

**Fig. S12. The virulence profile of eight FonR3-specific effector mutants in causing wilt disease of watermelon seedlings in preliminary greenhouse bioassays.** (A) The expression of eight putative FonR3-specific effectors in PDA medium (0 dpi) and during infection on cultivar G42 (1, 3, 6 dpi). (B, C, D) Disease progress of infected watermelon seedlings of PKR6, Sugarlee, and G42, respectively. For each treatment, 10 plants were used. Disease index was evaluated based on a 7-scale rating: 0 = asymptomatic, 1 = slight stunted growth and yellowing, 3 = stunted growth and yellowing, 5 = wilting, 7 = dead. Note: The FonR3Chr02G017230 gene was bold and given the name of *FonRSE1* in this study.

**Fig. S13. The *FonRSE1* is dispensable for vegetative growth and conidiation in the FonR3 isolate.** (A) Colony diameters of the WT FonR3, △*FonRSE1* deletion mutant, and complemented strain △*FonRSE1-C* were measured on PDA plates at 25 °C after 5 days. (B) Conidial production of the three indicated strains was counted by hemocytometer at 3 days post-inoculation within liquid potato dextrose broth (PDB) medium at 25°C in a 175-rpm shaker. (C) Under the same conditions, conidial morphology was photographed at 3 days post-inoculation. Scale bar = 50 μm. Mean and standard deviation (SD) of colony diameters and conidiation were estimated from three independent experiments. Different letters indicate Significant differences based on one-way ANOVA analysis followed by Duncan’s multiple range test (*p* = 0.05).

**Fig. S14. Two independent biological replicates of greenhouse bioassays for the functional study of *FonRSE1*.** For each treatment, eight to ten plants were tested. (A-B) Infected PKR6 plants at 28 dpi using 11-day-old seedlings for inoculations. (C-D) The corresponding disease index progress of infected PKR6 seedlings for four weeks. (E-F) The corresponding fresh weight measurements of the above-ground infected plants at 28 dpi. Different letters above the bars represent the significant difference between treatments using one-way ANOVA analysis followed by Duncan’s multiple range test (*p* = 0.05).

**Table S1. The information for putative FonR3-specific effectors.**

**Table S2. The list of primers used in the mutant constructions.**

**Table S3. The complete data sets for three independent greenhouse bioassays to functionally characterize the putative effector *FonRSE1* gene.**

