## Supplemental Figures for "Gap-free genomes and transcriptomes uncover race-specific effectors in watermelon wilt pathogen *Fusarium oxysporum* f. sp. *niveum*"

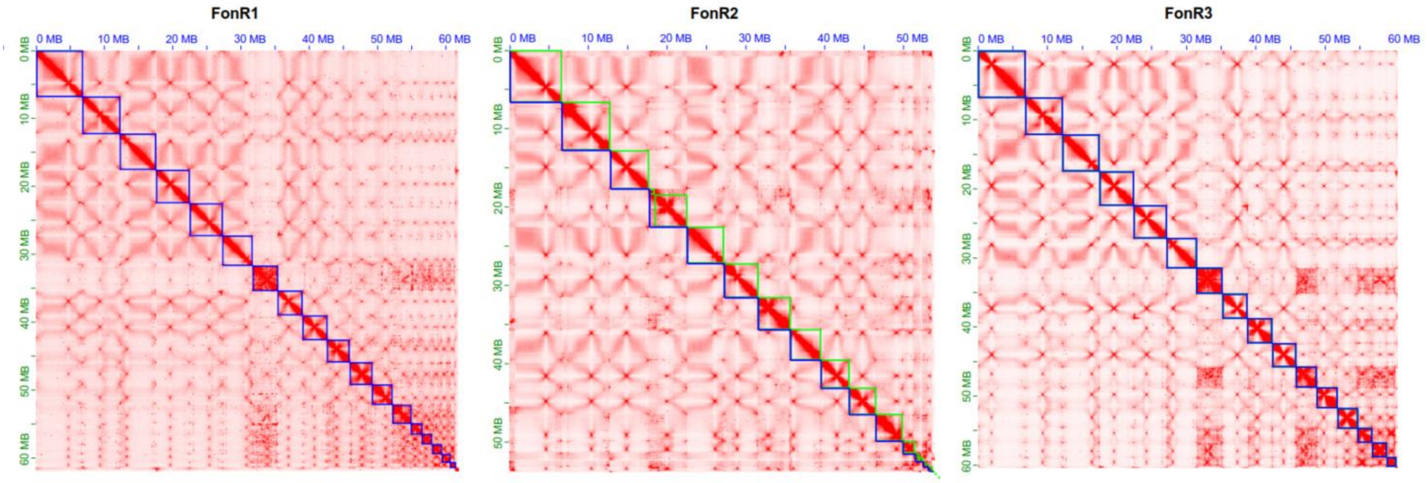


**Fig. S1. Hi-C contact maps of FonR1, FonR2, and FonR3 genome assemblies.**

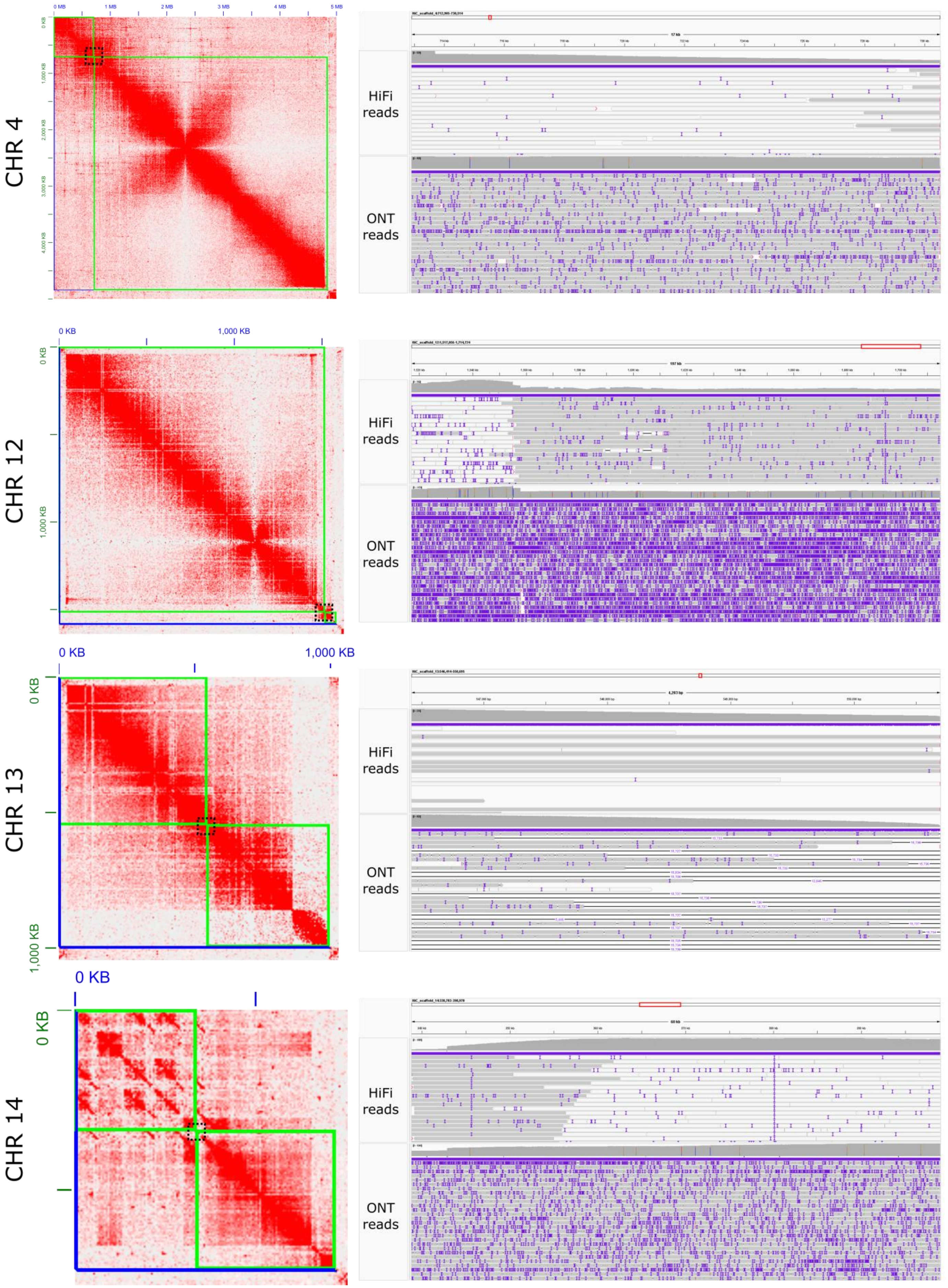


**Fig. S2. Confirmation of gap closing in FonR2 assembly.** On the left the chromosomes with gaps shown in black squares in the Hi-C contact maps, and on the right, IGV screenshots of HFi and ONT read mappings at the gap region.

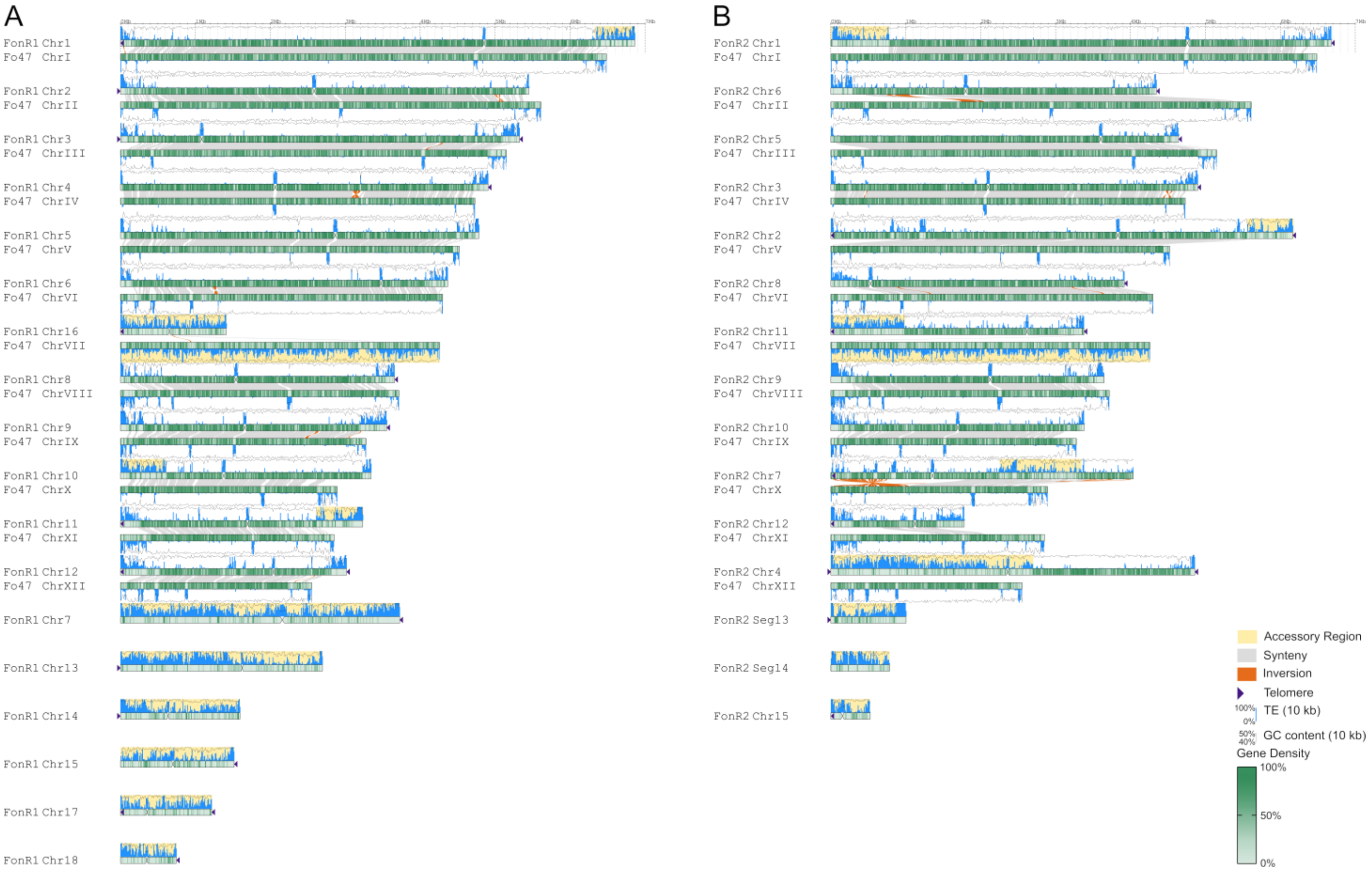


**Fig. S3. Genome alignment of FonR1, FonR2 and Fo47.** Ideograms of FonR1 (A) and FonR2 (B) were visualized using GenomeSyn[78]. Genomic features, including gene density, GC content, TE content, telomeres, centromeres, and genome synteny with Fo47 chromosomes are shown on the ideograms. The yellow background indicates the accessory regions.

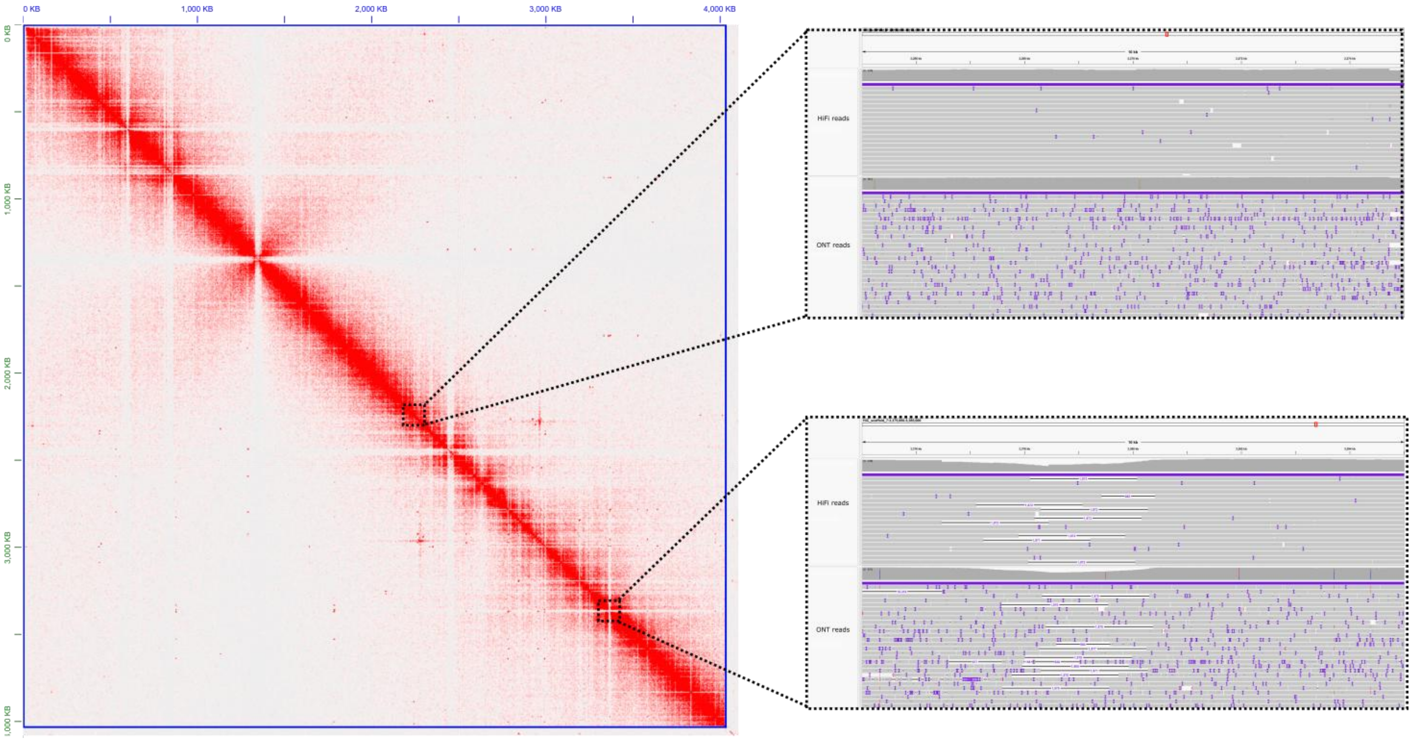


**Fig. S4. Confirmation of AR/CC border in FonR2.** Hi-C contact map showing Chr07 of FonR2 on the left and IGV screenshots with HiFi and ONT reads mappings on the AR/CC borders. AR: accessory regions. CC: core chromosome

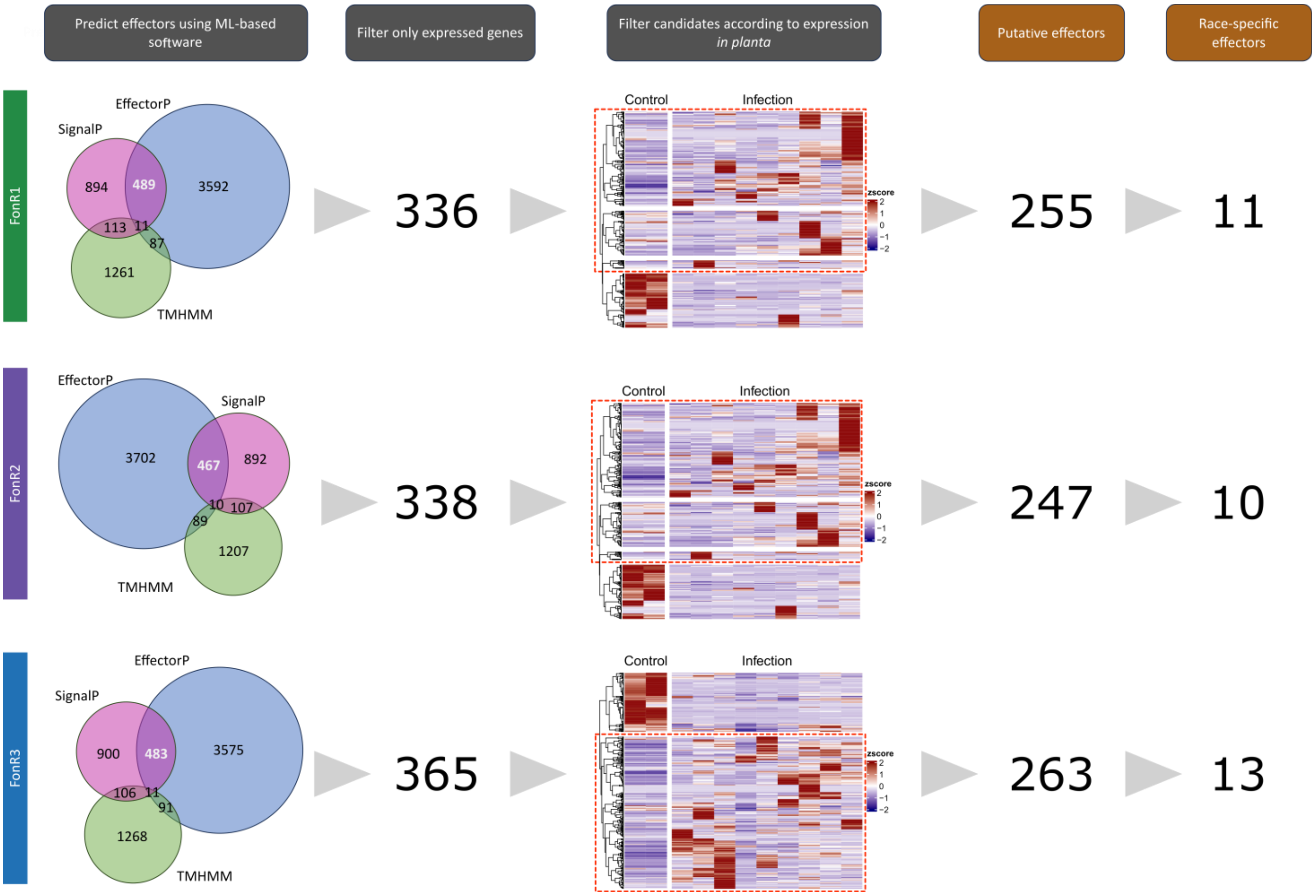


**Fig. S5. Fon effector prediction pipeline**. The effectors are first identified if they are predicted by SignalP[34] and EffectorP[33] but not including the transmembrane domain predicted by TMHMM[35]. After filtering, only the expressed genes in any of the selected conditions for transcriptome analysis, only genes that have increased expression in planta have been selected as putative effectors. Finally, race-specific effectors were identified with OrthoFinder[29].

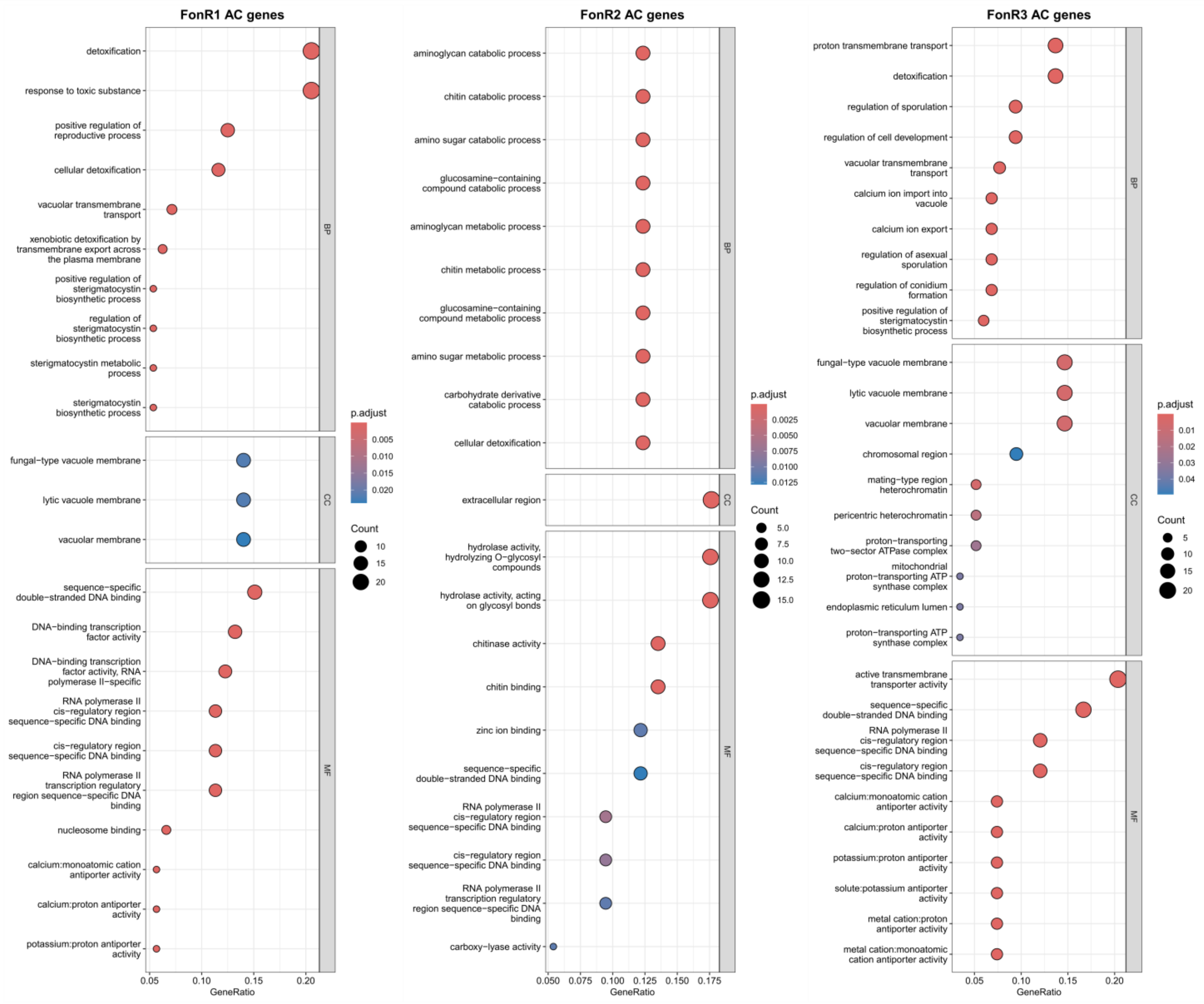


**Fig. S6. Gene ontology (GO) enrichment analysis of accessory chromosomes (AC) of FonR1 (left), FonR2 (middle), and FonR3 (right).** The color of bubbles is scaled by the adjusted p-values from enrichment analysis. The size of bubbles is scaled by the number of genes annotated with the specific GO terms. BP: biological process. CC: cellular component. MF: molecular function

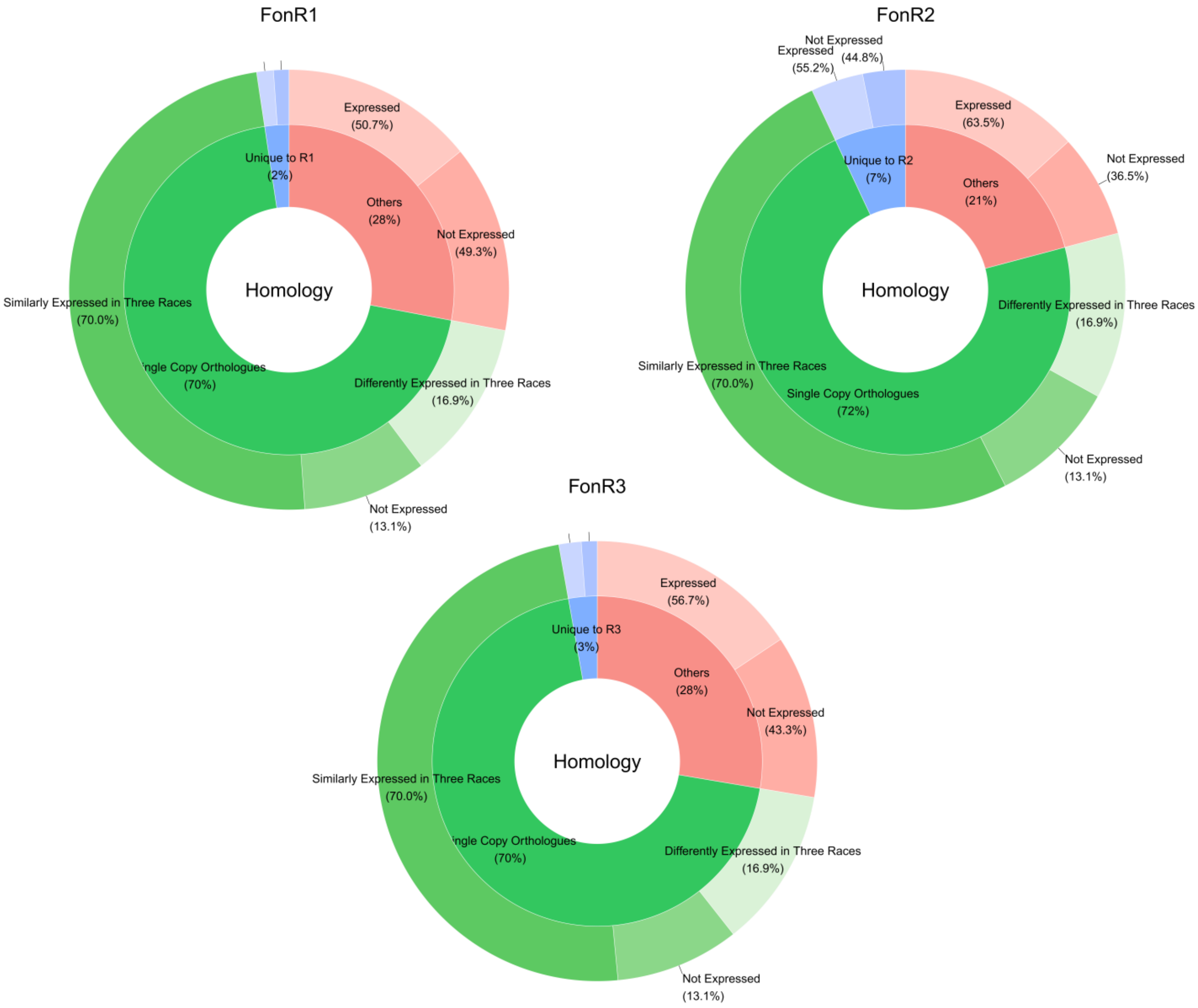


**Fig. S7. Donut plot summarizing the ratio of the ortholog gene family composition and gene expression conservation in annotated genes of three Fon races.**

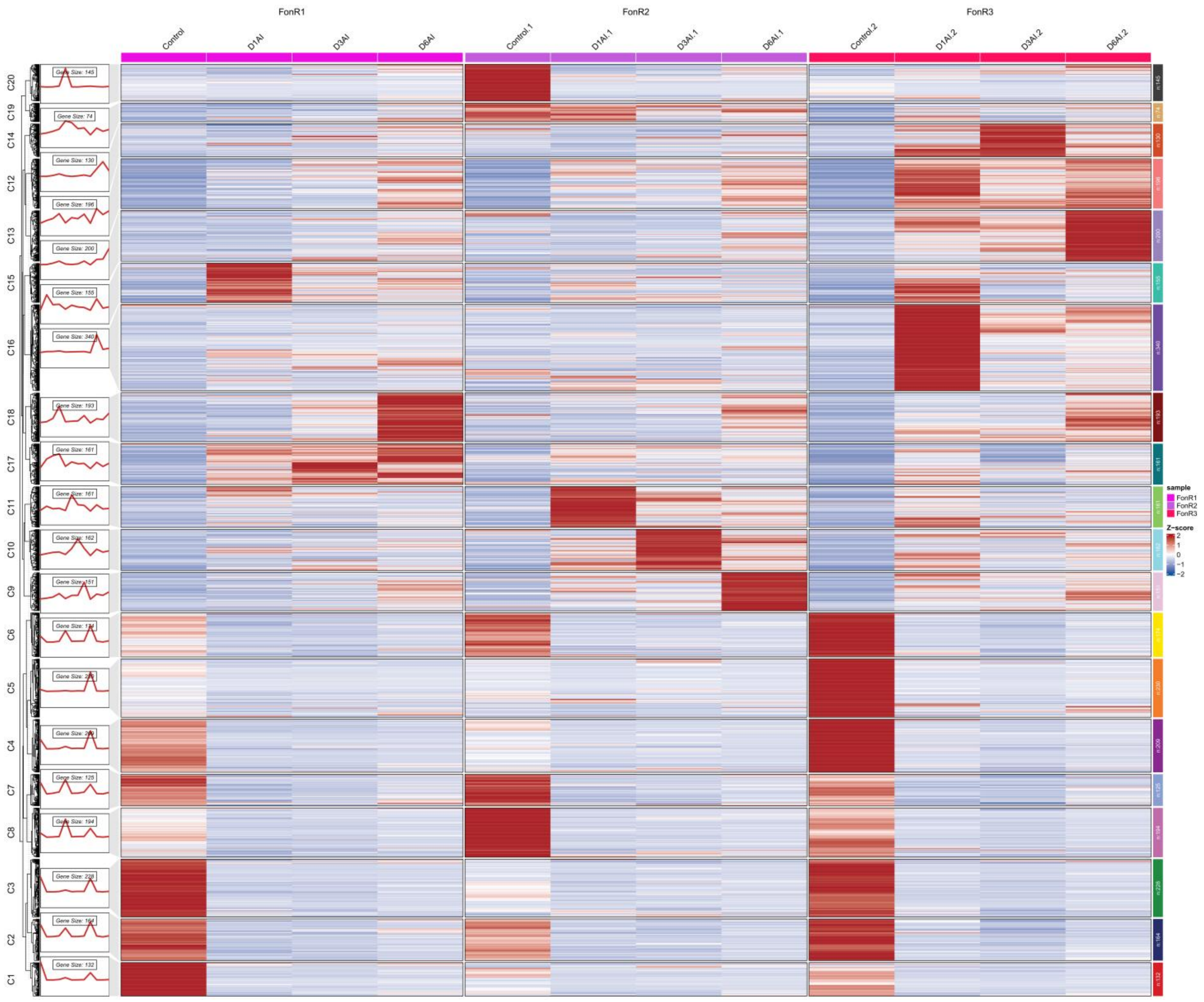


**Fig. S8. Heatmap showing the transcriptome clustering using K-means (k=20) clustering of single-copy ortholog expression of three races (FonR1, FonR2, and FonR3) at fungal culture and during infection of watermelon cultivar G42 at different time points (days) after inoculation (D1AI, D3AI, and D6AI).** Significantly enriched gene ontology terms are listed on the right side.

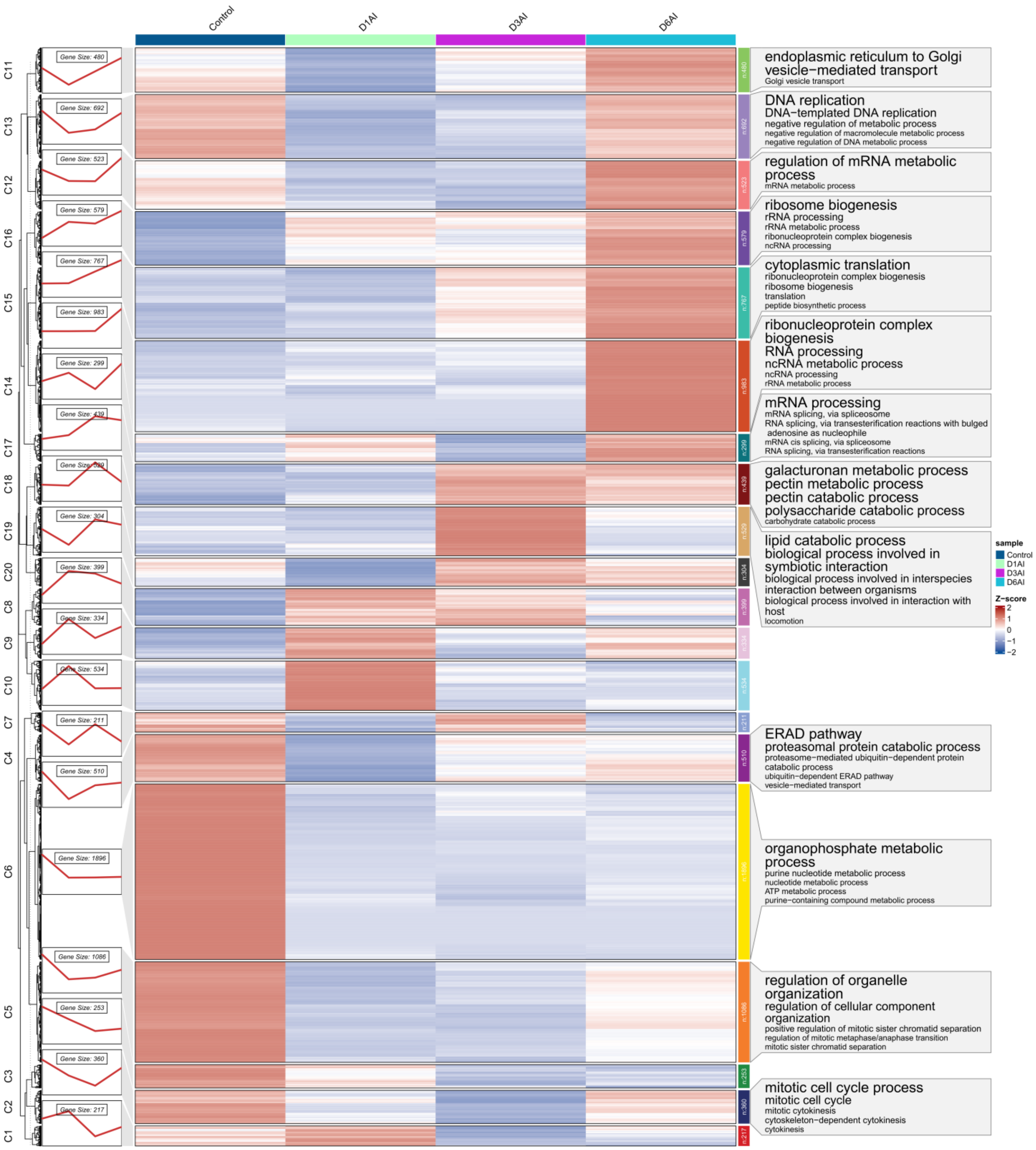


**Fig. S9. Heatmap showing the transcriptome clustering using K-means (k=20) clustering of FonR1 expression at fungal culture and during infection of watermelon cultivar G42 at different time points (days) after inoculation (D1AI, D3AI, and D6AI).** Significantly enriched gene ontology terms are listed on the right side.

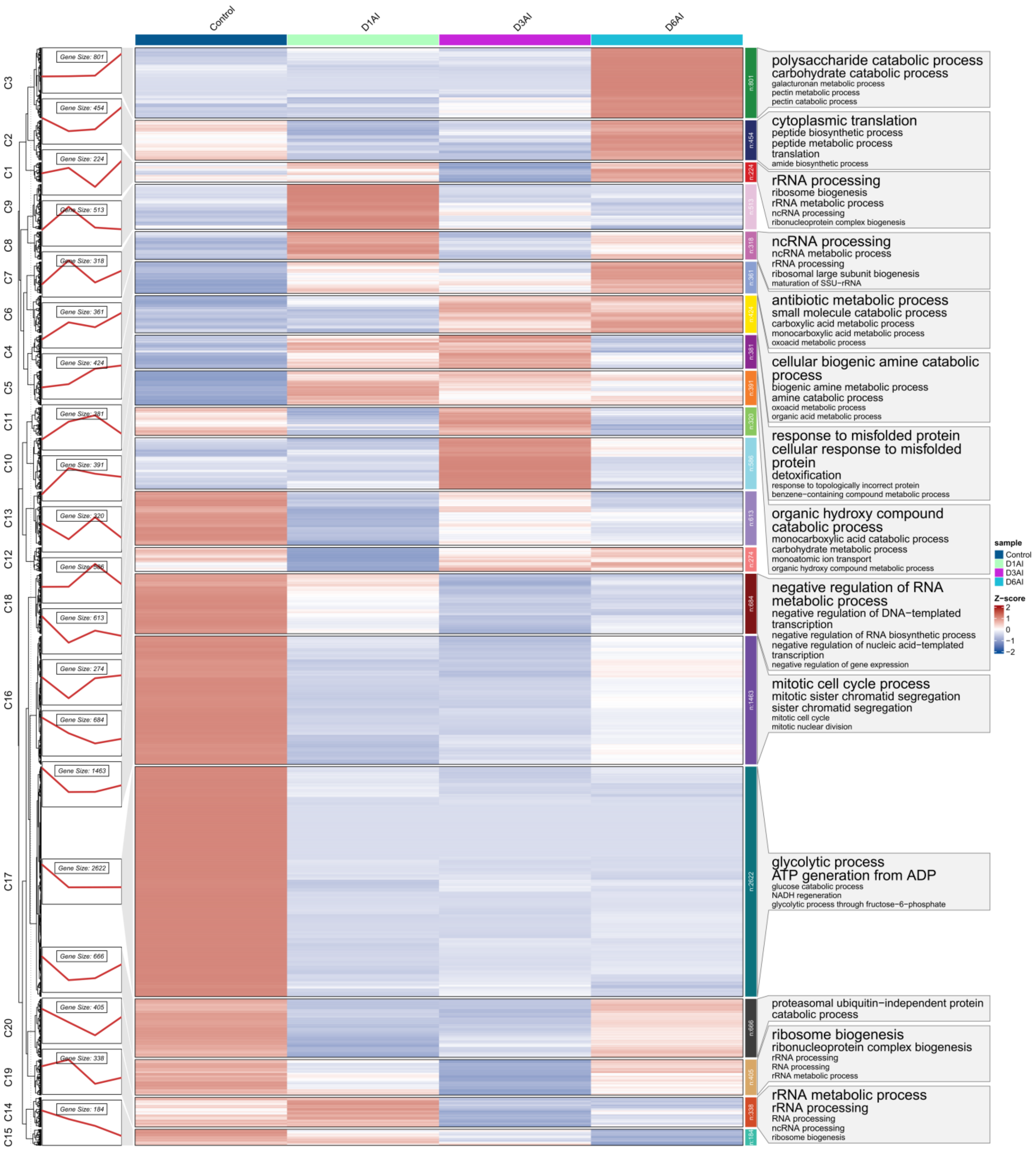


**Fig. S10. Heatmap showing the transcriptome clustering using K-means (k=20) clustering of FonR2 expression at fungal culture and during infection of watermelon cultivar G42 at different time points (days) after inoculation (D1AI, D3AI, and D6AI).** Significantly enriched gene ontology terms are listed on the right side.

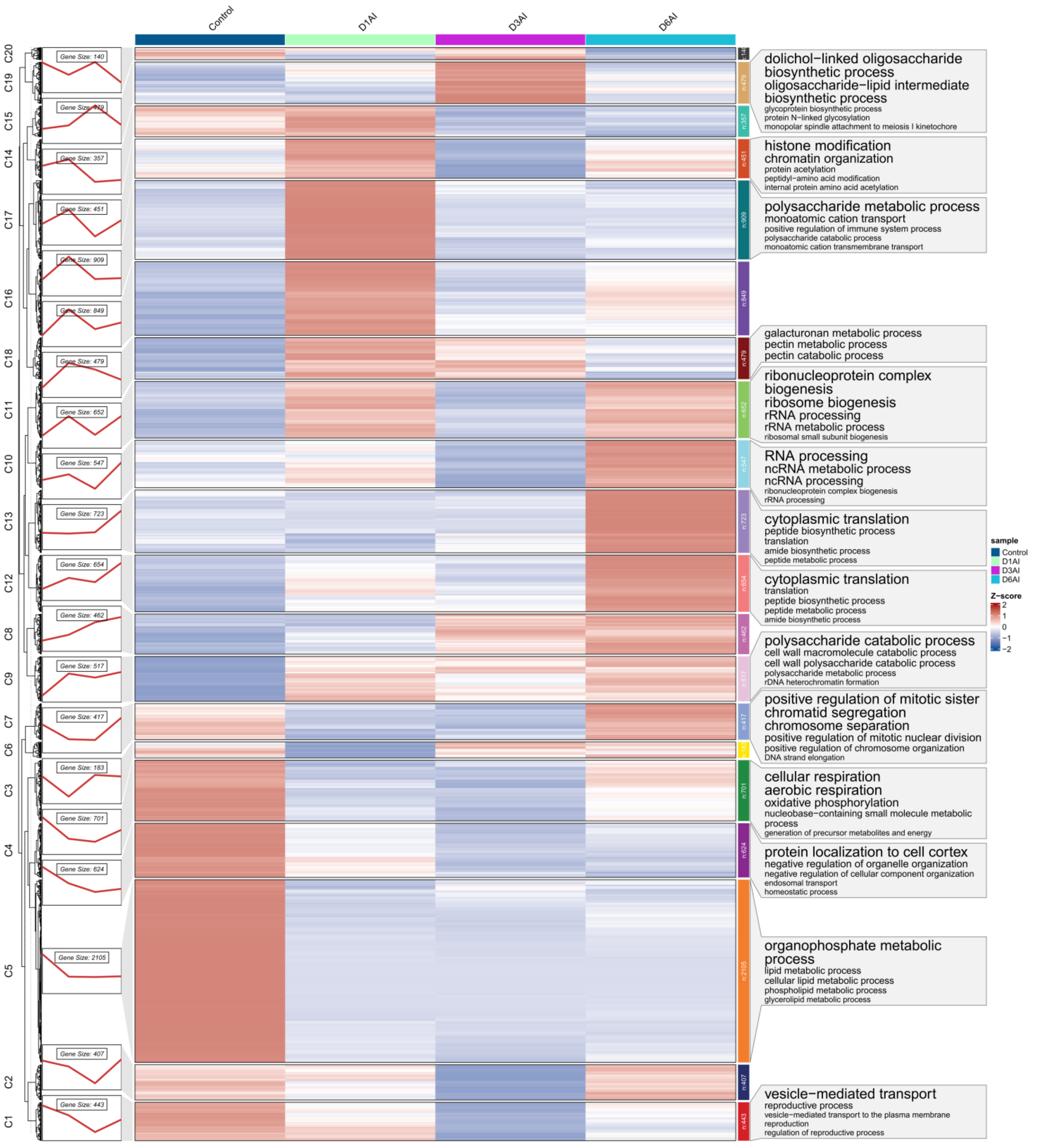


**Fig. S11. Heatmap showing the transcriptome clustering using K-means (k=20) clustering of FonR3 expression at fungal culture and during infection of watermelon cultivar G42 at different time points (days) after inoculation (D1AI, D3AI, and D6AI).** Significantly enriched gene ontology terms are listed on the right side.

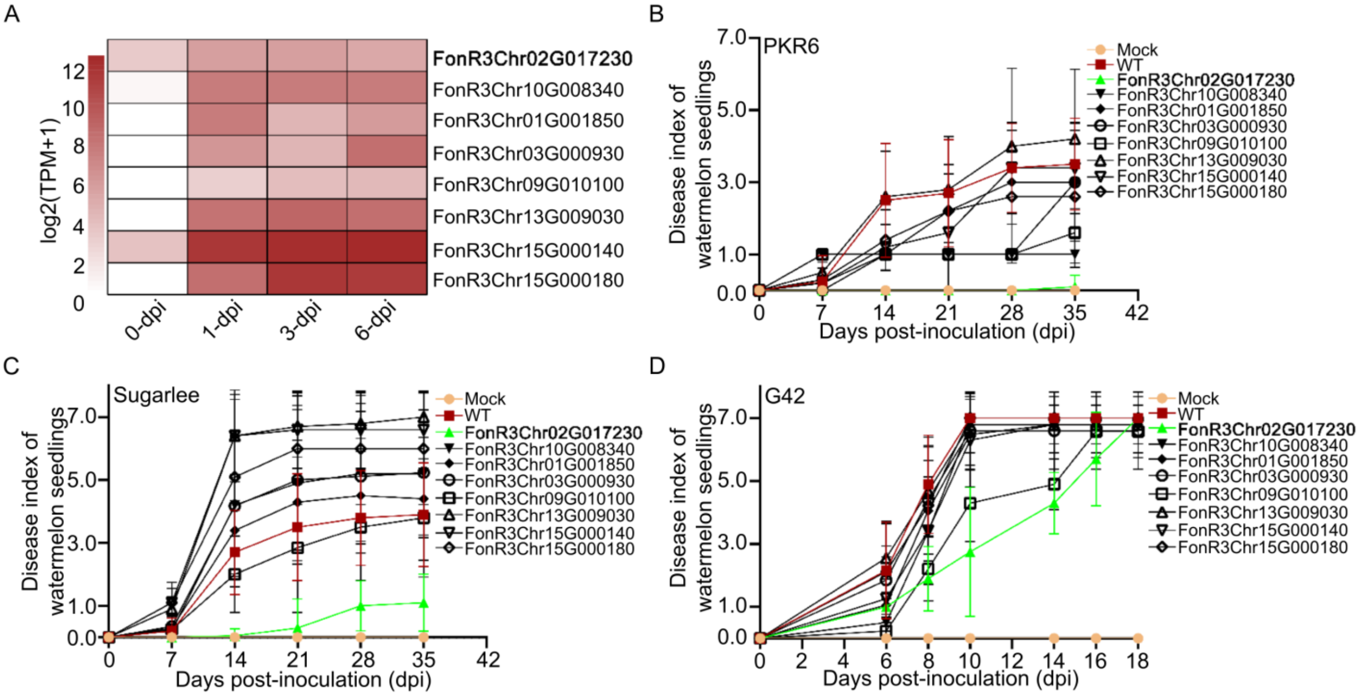


**Fig. S12. The virulence profile of eight FonR3-specific effector mutants in causing wilt disease of watermelon seedlings in preliminary greenhouse bioassays**. (A) The expression of eight putative FonR3-specific effectors in PDA medium (0 dpi) and during infection on cultivar G42 (1, 3, 6 dpi). (B, C, D) Disease progress of infected watermelon seedlings of PKR6, Sugarlee, and G42, respectively. For each treatment, 10 plants were used. Disease index was evaluated based on a 7-scale rating: 0 = asymptomatic, 1 = slight stunted growth and yellowing, 3 = stunted growth and yellowing, 5 = wilting, 7 = dead. Note: The FonR3Chr02G017230 gene was bold and given the name of *FonRSE1* in this study.

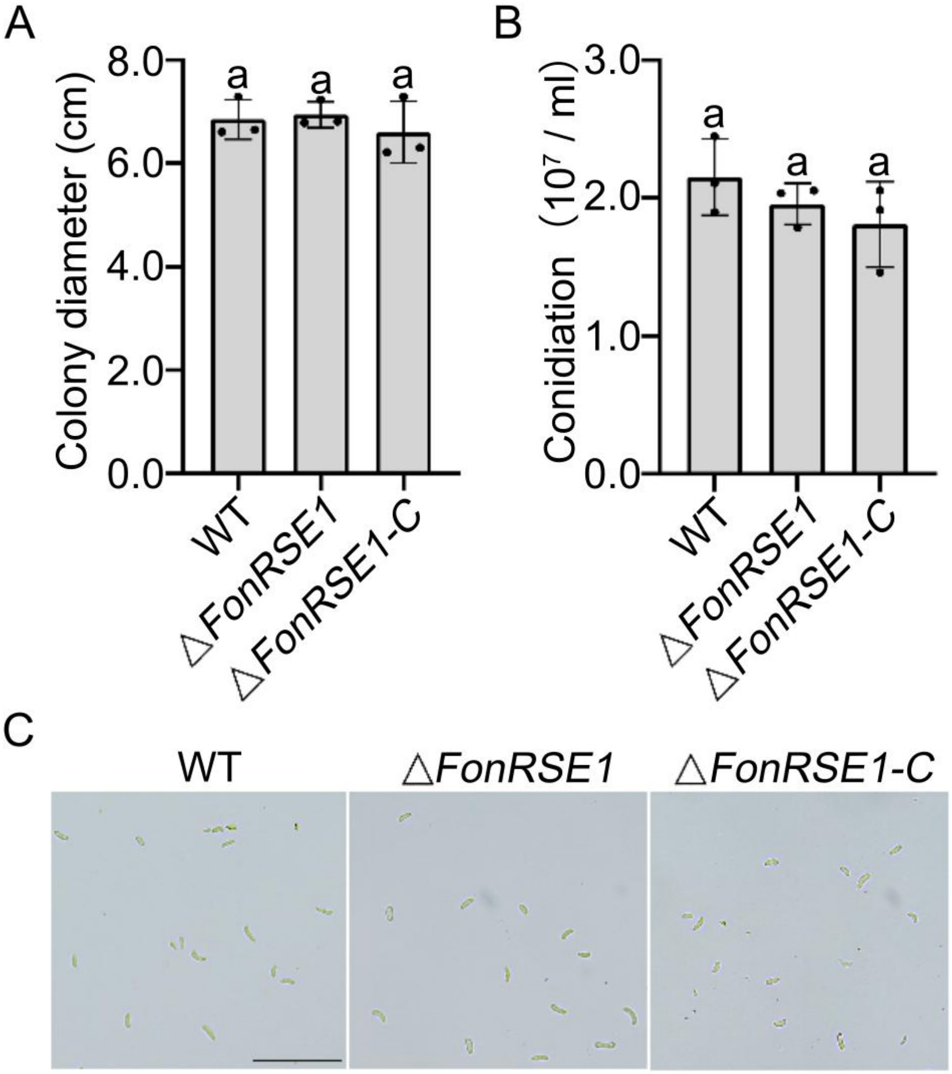


**Fig. S13. The *FonRSE1* is dispensable for vegetative growth and conidiation in the FonR3 isolate.** (A) Colony diameters of the WT FonR3, △*FonRSE1* deletion mutant, and complemented strain △*FonRSE1-C* were measured on PDA plates at 25 °C after 5 days. (B) Conidial production of the three indicated strains was counted by hemocytometer at 3 days post-inoculation within liquid potato dextrose broth (PDB) medium at 25°C in a 175-rpm shaker. (C) Under the same conditions, conidial morphology was photographed at 3 days post-inoculation. Scale bar = 50 μm. Mean and standard deviation (SD) of colony diameters and conidiation were estimated from three independent experiments. Different letters indicate significant differences based on one-way ANOVA analysis followed by Duncan's multiple range test ($p$ = 0.05).

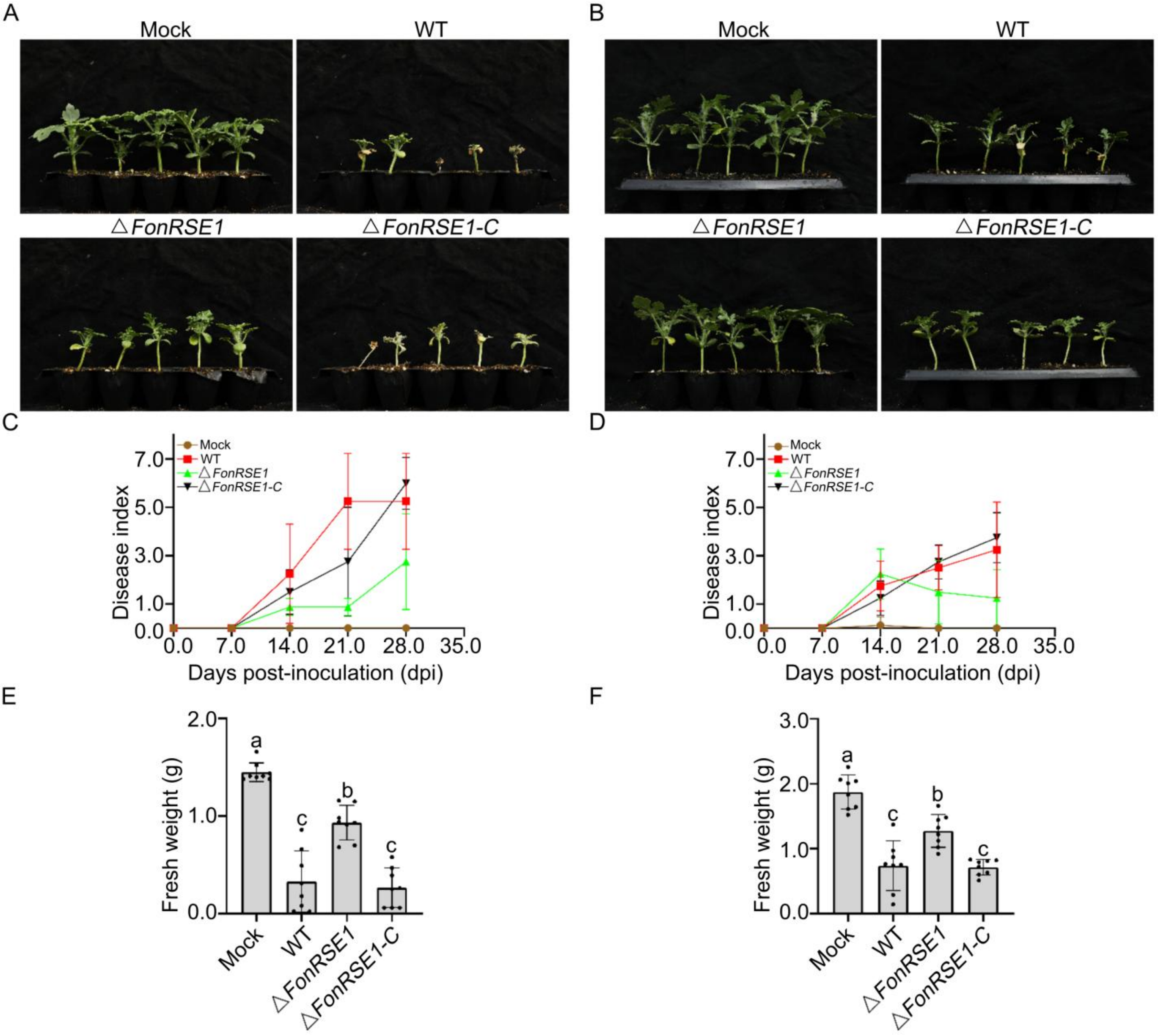


**Fig. S14. Two independent biological replicates of greenhouse bioassays for the functional study of FonRSE1.** For each treatment, eight to ten plants were tested. (A-B) Infected PKR6 plants at 28 dpi using 11-day-old seedlings for inoculations. (C-D) The corresponding disease index progress of infected PKR6 seedlings for four weeks. (E-F) The corresponding fresh weight measurements of the above-ground infected plants at 28 dpi. Different letters above the bars represent the significant difference between treatments using one-way ANOVA analysis followed by Duncan's multiple range test ($p = 0.05$).
